# Bridging Gaps in Antibody Responses and Animal Welfare: Assessing Blood Collection Methods and Vaginal Immunity in Mice Immunized with Intranasal Gonococcal Vaccines

**DOI:** 10.1101/2025.02.23.639724

**Authors:** Abhishek Chanda, Yujuan Song, Junaid Nazir, Chenwei Lin, Alicia Cheng, Jennifer Sargent, Aleksandra E. Sikora

## Abstract

Assessing antibody titers and functional responses is essential for evaluating vaccine efficacy, yet the impact of blood collection methods on these immunological assessments remains unclear. Retro-orbital (RO) blood collection is commonly used but significant complications can occur. Increasingly, investigators have adopted alternative blood collection approaches, such as saphenous vein (SV) sampling to improve laboratory animal welfare. This study compared RO and SV sampling in the development of a *Neisseria gonorrhoeae* (Ng) vaccine, evaluating Adhesin Complex Protein (ACP) and multiple transferable resistance (Mtr) E protein (MtrE) as antigen candidates. Epitope mapping revealed that ACP and MtrE possess multiple, highly accessible B-cell and T-cell epitope clusters, reinforcing their immunological potential. Following intranasal immunization with rACP, rACP+CpG, and rMtrE+CpG, we assessed the specificity, magnitude, kinetics, and functional quality of immune responses elicited by the immunization regimens. Out of 45 comparisons, only eight significant differences were detected in antibody titers, while the human serum bactericidal assays revealed no differences between RO and SV in antigen-immunized groups. However, antibodies elicited by rACP alone or ACP+CpG in SV samples restored 30.05% and 75.2% of human lysozyme hydrolytic activity compared to 19.3 and 59.9 % in RO, respectively suggesting that SV sampling may be more reliable for assessing functional antibody responses. Beyond its immunological advantages, SV sampling reduces stress, minimizes ocular trauma, and improves animal welfare, making it a viable alternative to RO collection. Given its widespread use in vaccine research, standardizing SV sampling could improve data reliability, ethical compliance, and translational relevance in preclinical studies.

## INTRODUCTION

In preclinical vaccine studies, assessment of antibody titers and functional responses are pivotal endpoints, serving as critical indicators of vaccine efficacy and correlates of protection. However, an often-overlooked factor in these assessments is how blood collection methods may influence experimental outcomes. While retro-orbital (RO) bleeding has been widely used for its efficiency in yielding large blood volumes, it carries substantial risks, including inflammation, hemorrhage, hematomas, fractures, and potential blindness [1–5]. These concerns have led to increasing adoption of less invasive methods, such as saphenous vein (SV) sampling, which minimizes animal distress and aligns with ethical principles of the 3Rs (Replacement, Reduction, and Refinement) [1, 2, 6]. However, it remains unclear whether blood collection techniques introduce variability in antibody levels or functional immune responses.

To address this gap, we examined blood sampling methods in the context of developing a vaccine against *Neisseria gonorrhoeae* (Ng, gonococcus), a highly prevalent and increasingly drug-resistant pathogen responsible for gonorrhea. Ng is a Gram-negative bacterium with strict host tropism for humans, colonizing the mucosal surfaces of the eye, oropharynx, rectum, male urethra, and female ecto- and endocervix. Transmission of Ng occurs through sexual contact and during prenatal exposure to mother’s infected cervix [7, 8]. Activation of innate immune responses at the colonization sites are largely responsible for the pathological damage, and gonorrhoea, if undiagnosed or inappropriately treated, may have severe consequences on reproductive and neonatal health. The sequela can include salpingitis, pelvic inflammatory disease, ectopic pregnancy, infertility, urethral strictures, and rarely disseminated disease. Gonococcal infection among neonates usually manifests as acute illness several days after birth with most common ophthalmia neonatorum. Gonorrhoea remains an important public health issue globally with an estimated 84.2 million of new cases among adults aged 25-49 years in 2020 [9]. A single monotherapy of intramuscular ceftriaxone is currently recommended for treatment of uncomplicated urogenital, anorectal and pharyngal Ng infections [10]. Ng antibiotic resistance to many antibiotics, however, increased rapidly in recent years and a rise in ceftriaxone-resistance poses a formidable challenge to gonorrhoea control [9, 11]. People with Ng infection are at higher risk of acquisition and transmission of human immunodeficiency virus, which further necessitate the need to develop safe and effective gonococcal vaccines [8, 9].

Current vaccine strategies focus on evaluating immune responses induced by a variety of Ng vaccine antigens formulated with different adjuvants and delivered via subcutaneous, intraperitoneal, intravaginal, intramuscular, or intranasal administration routes [9]. Among the promising gonococcal vaccine candidates are Ng Adhesin Complex Protein (Ng-ACP) and the multiple transferable resistance (Mtr) E protein due to their conservation, pivotal functions, antigenicity and ability to induce bactericidal antibodies [12–17].

Ng-ACP was originally identified in a closely related to Ng, *N. meningitidis* (Nm), and it is a dual function surface-exposed protein that acts as adhesin and an inhibitor of human lysozyme in Ng, Nm and commensal *Neisseria* species [14–16]. Although Ng-ACP and Nm-ACP do not contain the conserved sequence motifs required for lysozyme recognition and binding, their three dimensional structures closely resemble those of MliC/PliC members of lysozyme inhibitors [15, 18]. Both mice and rabbits develop functional blocking antibodies that preclude ACP from inhibiting the lytic activity of human lysozyme [18, 19].

MtrE is a homotrimeric outer membrane β-barrel protein that serves as a channel for MtrCDE, FarAB and MacAB efflux pumps that export antibiotics, fatty acids and antimicrobial products of the innate defence [20–23]. Ng mutants that are deficient in MtrE or MtrA, an activator of *mtrECD* operon, showed significantly reduced fitness in the lower genital tract of female mouse [24, 25]. Additionally, MtrE enhances Ng survival in neutrophil extracellular traps, showcasing its role in immune evasion [26]. While MtrE is pivotal for in vivo fitness in gonococcal murine infection model, its necessity in human infections remains inconsistent across studies. However, MtrE’s role in bacterial defense mechanisms underscores its potential as a vaccine target [27].

In this study, we further evaluated MtrE and ACP as potential gonococcal vaccine antigens by identifying their predicted B cell and T cell epitopes using available crystal structures. We then examined whether antibody levels and functional immune responses differed between RO and SV blood collection sites in mice following intranasal immunization with recombinant, mature ACP (rACP), rACP combined with the TLR9 agonist ODN 2395 class C cytosine–phosphate–guanine (hereafter CpG), and rMtrE adjuvanted with CpG. By comparing these two blood sampling methods, we aimed to assess whether a shift toward less invasive techniques could enhance research reliability and animal welfare without compromising immunological assessments. Despite the elusive nature of immune correlate of protection for gonococcal vaccines [9], antibody titers remain key indicators of vaccines efficacy, offering valuable insights into immune responses. Beyond gonococcal vaccine research, this study contributes to the broader movement toward refining experimental methodologies, ensuring vaccine development remains scientifically rigorous while aligning with best practices in laboratory animal care. As vaccinology advances, integrating methodological improvements with immunological research will be crucial for accelerating the development of next-generation vaccines that address pressing global health challenges.

## RESULTS

### B cell and T cell epitopes prediction and mapping for ACP and MtrE antigens

ACP is localized on the surface of Ng and Nm, as part of their outer membrane, facing the extracellular environment. This has been confirmed by fluorescence-activated cell sorter analysis, which demonstrated surface expression of ACP and its accessibility to antibodies [15, 16, 18]. Structurally, ACP adopts an eight-stranded β-barrel, where the β1-β4 sheet is predominantly basic and faces outward and the loop regions between β4 and β5 and between β8 and β1 are surface-exposed [15, 18]. In contrast to ACP, most of MtrE is embedded in the Ng cell envelope. MtrE forms a transmembrane β-barrel in the outer membrane that extends into the periplasm. The extracellular loops (L1-L3) of the β-barrel are oriented outward, making them accessible on the bacterial surface [22]. To further evaluate ACP and MtrE as potential gonococcal antigens, computational tools were applied to the available crystal structures of both proteins. These analyses included the prediction of B-cell epitopes, which are critical for antibody recognition, as well as MHC class I and MHC class II epitopes, which are essential for T-cell mediated immune responses (**Fig. 1**, **Tables 1**-**2**). For ACP, a total of four B-cell epitopes (**Fig. 1A-D**), ranging from 5 to 23 amino acids in length, were identified as high-affinity binders with surface exposure scores exceeding 0.6. These epitopes are predominantly located within the N-terminal domain (**Table 1**) and are mapped to surface-accessible regions (**Fig. 1A-D**). The predicted MHC I (10 peptides, **Fig. 1E**, **Table 1**) and MHC II epitopes (5 peptides, **Fig. 1F**, **Table 1**) are strategically distributed across ACP and involve peptides essential for T-cell activation. The B-cell epitopes (e.g., chain A: V41, N43, G44, K45) and MHC-binding epitopes (e.g., GTDNPTVAK for MHC-I and YVCQQGKKV for MHC-II) are located within regions corresponding to the mature ACP. These epitopes map to regions of the protein that can effectively stimulate both CD8+ and CD4+ T-cells, fostering robust cellular immunity. Specifically, B-cell epitopes (e.g., Chain A: V41, G44, K45) map to surface-exposed loops, optimizing their immunogenicity, MHC I epitopes (e.g., GTDNPTVAK) align with regions that bind to HLA-A alleles, critical for CD8+ T-cell response. Whereas MHC II epitopes (e.g., YVCQQGKKV) are well positioned to facilitate interaction with HLA-DR alleles for CD4+ T-cell activation.

**Figure 1.**
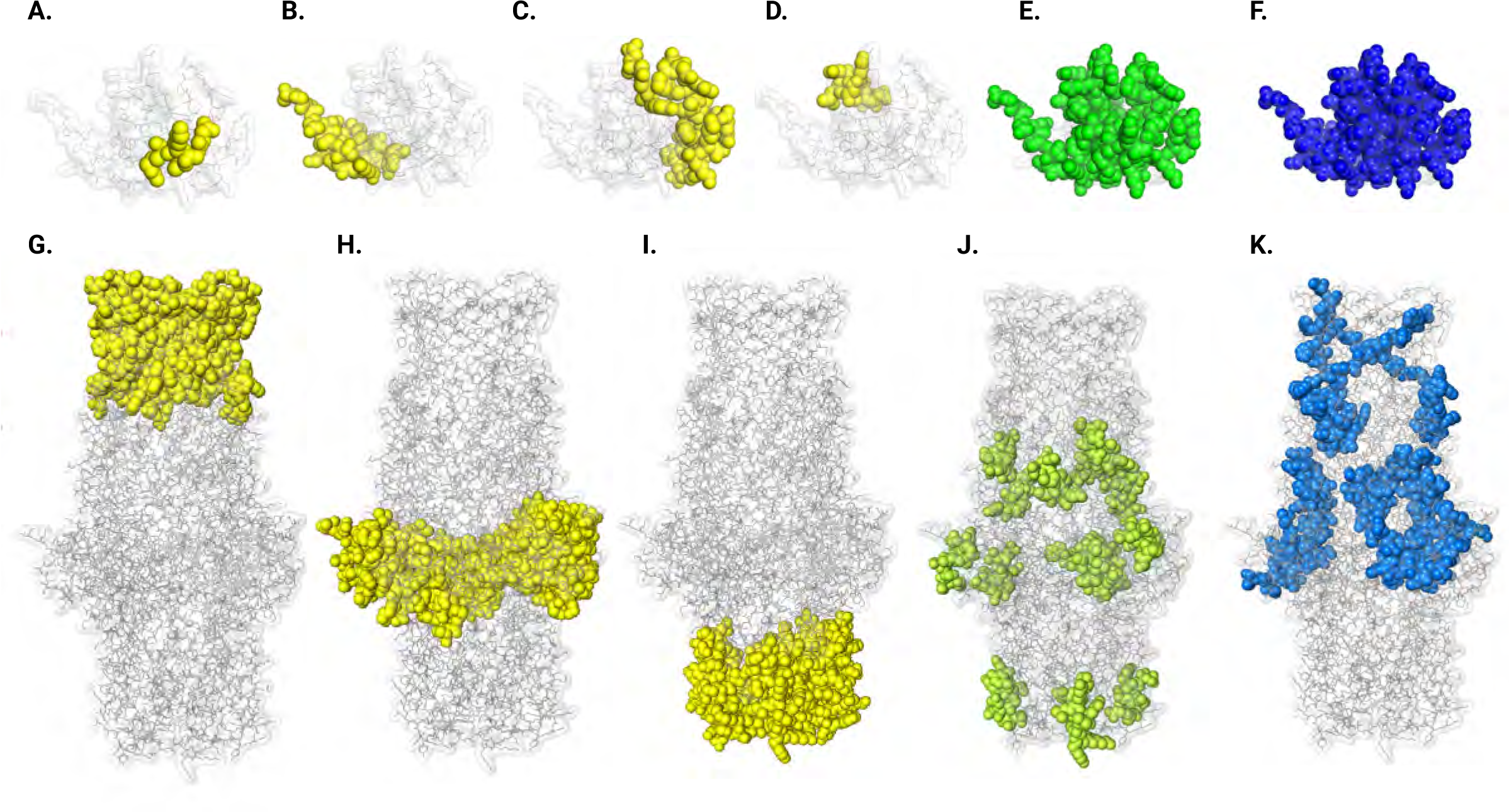
Visualization of predicted B Cell, MHC I, and MHC II epitopes on ACP (A-F) and MtrE (G-K) 3D structures. Panels **A**–**D** highlight B cell epitopes (yellow) mapped on the ACP structure, revealing surface-accessible regions suitable for antibody binding. Panels **E** and **F** show MHC I (green) and MHC II (blue) epitopes on ACP, respectively, identifying immunogenic peptides potentially recognized by T cells. Panels **G**–**I** depict B cell epitopes (yellow) distributed across the MtrE structure. Panels **J** and **K** represent MHC I (green) and MHC II (blue) epitopes on MtrE, emphasizing regions with affinity for T cell activation.

**Table 1.** Epitope clusters identified on ACP crystal structure with predicted B-cell emphasizing, MHC-I and MHC-II binding sites.

| Source | Length (AA) | Allele | Score |
| --- | --- | --- | --- |
| **B-cell Epitopes (Peptides)** |  |  |  |
| Chain A: V41, N43, G44, K45, R46 | 5 | NA | 0.767 |
| Chain A: A1, G2, T3, D4, N5, P6, T7, V8, A9, K10, N30, K31, Q32, G33, L34, D77, S78, K79, Y81 | 19 | NA | 0.682 |
| Chain A: V16, C17, Q18, Q19, G20, K21, K22, E66, G67, G68, Y69, A90, P91, D92, N93, Q94, I95, V96, K98, D99, S101, P102, R103 | 23 | NA | 0.671 |
| Chain A: N52, L53, D54, K55, S56, D57, N58, M59 | 8 | NA | 0.655 |
| **MHC I (Peptides)** |  |  |  |
| GTDNPTVAK | 9 | HLA-A*11:01, HLA-A*03:01 | 0.927 |
| TVAKKTVSY | 9 | HLA-A*26:01, HLA-B*15:01, HLA-A*30:02, HLA-B*35:01, HLA-A*68:01 | 0.946 |
| QQGKKVKVTY | 10 | HLA-B*15:01 | 0.886 |
| YASAVINGK | 9 | HLA-A*68:01 | 0.910 |
| RVQMPINLDK | 10 | HLA-A*03:01, HLA-A*11:01 | 0.858 |
| DKSDNMDTFY | 10 | HLA-A*01:01, HLA-A*26:01 | 0.972 |
| GAMDSKSYR | 9 | HLA-A*68:01, HLA-A*31:01, HLA-A*11:01 | 0.764 |
| AMDSKSYRK | 9 | HLA-A*03:01, HLA-A*11:01 | 0.799 |
| SYRKQPIMI | 9 | HLA-B*57:01, HLA-A*24:02 | 0.711 |
| ITAPDNQIVF | 10 | HLA-B*57:01, HLA-B*58:01, HLA-B*35:01 | 0.803 |
| **MHC II (Peptides)** |  |  |  |
| YVCQQGKKV | 9 | HLA-DRB1*01:01 | 0.923 |
| LTTYASAVI | 9 | HLA-DRB1*15:01 | 0.945 |
| INLDKSDNM | 9 | HLA-DRB1*03:01 | 0.745 |
| YVLSTGAMD | 9 | HLA-DRB1*04:05 | 0.917 |
|  |  | HLA-DRB1*07:01 |  |
|  |  | HLA-DRB1*09:01 |  |
| IMITAPDNQ | 9 | HLA-DRB4*01:01 | 0.828 |

For the MtrE homotrimer, predictions highlight a broad distribution of B-cell epitopes across the extensive surface areas of the protein, including both surface-exposed loops and regions embedded in the outer membrane that extend into the periplasmic space (**Fig. 1G-I**, **Table 2**). Notably, clusters of epitopes comprising residues G889–G1215, spanning 267 amino acids, are prominently located within the surface-exposed loops and outer membrane region (**Fig. 1G**, **Table 2**). The predicted MHC I and MHC II epitopes for MtrE, comprising 10 and 8 peptides respectively, are similarly well-positioned to engage T-cell responses (**Fig. 1J-K**, **Table 2**). The MHC I epitopes (e.g., AVLSNEIYR) map to periplasmic-localized regions of MtrE that can be potentially recognized by diverse HLA alleles, maximizing CD8+ T-cell engagement (**Fig. 1J**). The MHC II epitopes (e.g., FNQDSYVSS), identified in periplasmic and outer membrane parts of MtrE, target HLA-DRB1 alleles (**Fig. 1K**), ensuring effective helper T-cell activation for a sustained immune response.

**Table 2.** Epitope clusters identified on MtrE crystal structure with predicted B-cell, MHC-I and MHC-II binding sites.

| Source | Length<br>(AA) | Allele | Score |
| --- | --- | --- | --- |
| **B-cell Epitopes (Peptides)** |  |  |  |
| **Chains: A, B, C** | **267** | **NA** | **0.769** |
| G889, K890, I892, P893, Q894, L966-L967, P968, T969, L970, A971-A972, N973, A974, N975, G976, S977, R978, Q979, G980, S981, L982, S983, G984-G985, N986, V987, S988-S990, Y991, N992, V993, G994, L995, G996, A997-A998, S999, Y1000, E1001, L1002, D1003, L1004, F1005, G1006, R1009, F1173-F1174, P1175, S1176, I1177, L1179, T1180, G1181, S1182, V1183, G1184, T1185, G1186, S1187, V1188, E1189, L1190, G1191-G1192, L1193, F1194, K1195, S1196, G1197, T1198, G1199, V1200, W1201, A1202, F1203, A1204, P1205, S1206, I1207, T1208, L1209, P1210, I1211, F1212, T1213, W1214, G1215 |  |  |  |
| **Chains: A, C** | **68** | **NA** | **0.642** |
| V900, E901, V902, A903, E904, T905, F906, N908, D909, T910, S911, V912, S913-S914, V915, R916, A917, V918, D919, L920, G921, W922, H923, D924, A927, D928, P929, R930, Q932, I1114, N1115, R1116, P1117, I1118, P1119, E1120, D1121, L1122, P1123, A1124, G1125, L1126, P1127, D1129, K1130, Q1131, F1133, K1136, L1137, P1138, A1139, G1140, L1141, S1142-S1143, E1144, D1148, L1323, G1324-G1326, L1327, K1328, R1329, D1330, T1331, Q1332, T1333 |  |  |  |
| **Chains: A, B, C** | **183** | **NA** | **0.767** |
| S1052, Q1055, R1056, V1057, L1058, K1059, T1060, R1061, E1062-E1063, T1064, Y1065, K1066, L1067, S1068, E1069, L1070, R1071, Y1072, K1073, A1074, G1075, V1076, I1077, S1078, A1079, V1080, A1081, L1082, R1083, Q1084-Q1085, E1086, I1089, R1268, A1269, S1270, K1271, E1272, A1273, L1274, R1275, L1276, V1277, G1278, L1279, R1280, Y1281, K1282, H1283, G1284, V1285, S1286, G1287, A1288, L1289, D1290, L1291, L1292, A1294, E1295 |  |  |  |
| **MHC I (Peptides)** |  |  |  |
| SSVRAVDLGW | 9 | HLA-B*57:01, HLA-B*58:01 | 0.967 |
| AVDLGWHDY | 9 | HLA-A*01:01 | 0.886 |
| KLIDIALER | 10 | HLA-A*31:01, HLA-B*08:01 | 0.886 |
| AVLNSEIYR | 9 | HLA-B*35:01, HLA-A*11:01, HLA-A*31:01 | 0.890 |
| YMIERNNLL | 10 | HLA-A*02:01, HLA-A*02:03 | 0.938 |
| SSSEAALQGY | 10 | HLA-A*01:01, HLA-B*44:03 | 0.942 |
| ALQGYFASV | 9 | HLA-A*02:03, HLA-A*02:01 | 0.914 |
| KTREETYNA | 9 | HLA-B*15:01 | 0.912 |
| ETYNAVRIAV | 9 | HLA-A*68:02 | 0.881 |
| PSITLPIFTW | 10 | HLA-B*57:01, HLA-B*58:01 | 0.931 |
| **MHC II (Peptides)** |  |  |  |
| FQNDTSVSS | 9 | HLA-DRB1*04:01 | 0.973 |
| YMIERNNLL | 9 | HLA-DRB1*04:01, HLA-DRB1*01:01 | 0.965 |
| YFASVANRD | 9 | HLA-DRB1*04:05, HLA-DRB1*09:01, HLA-DRB1*04:01 | 0.965 |
| FFPSIRLTG | 9 | HLA-DRB1*11:01, HLA-DRB1*08:02 | 0.957 |
| WAFAPSITL | 9 | HLA-DRB1*09:01, HLA-DRB1*07:01 | 0.878 |
| FAPSITLPI | 9 | HLA-DRB1*09:01 | 0.873 |
| IVAYESAVQ | 9 | HLA-DRB1*15:01 | 0.946 |
| YKALGGGLK | 9 | HLA-DRB5*01:01 | 0.855 |

Together, the epitope mapping demonstrated that ACP and MtrE possess multiple, highly accessible epitope clusters with broad immunogenic potential. These findings, combined with their functional roles in bacterial virulence—ACP’s involvement in adhesion and immune evasion, and MtrE’s contribution to antimicrobial resistance through the MtrCDE efflux pump—strongly support the appraisal of both proteins as promising gonococcal vaccine candidates capable of eliciting protective immune responses.

### Preparation of rACP and rMtrE vaccines and immunization experiments

Signal peptides, which direct proteins to the bacterial cell envelope or extracellular milieu, are cleaved off during protein maturation, leaving only the functional regions exposed. Including these signal peptide sequences in vaccine antigens could trigger immune responses against irrelevant regions, thereby diminishing vaccine efficacy. Computational epitope mapping of ACP and MtrE revealed significant immunogenic clusters on their mature forms, particularly in surface-exposed loops and regions critical for bacterial virulence (**Fig. 1**, **Tables 1**–**2**). To focus the immune response on these biologically relevant epitopes, we engineered a recombinant version of ACP lacking its native signal peptide (amino acid residues 1–21, **Fig. 2A**) and used a previously optimized construct to express mature MtrE (**Fig. 2B**, [28]). This strategy ensured accurate representation of the immunologically active forms of these proteins, which is crucial for effective vaccine development.

**Figure 2.**
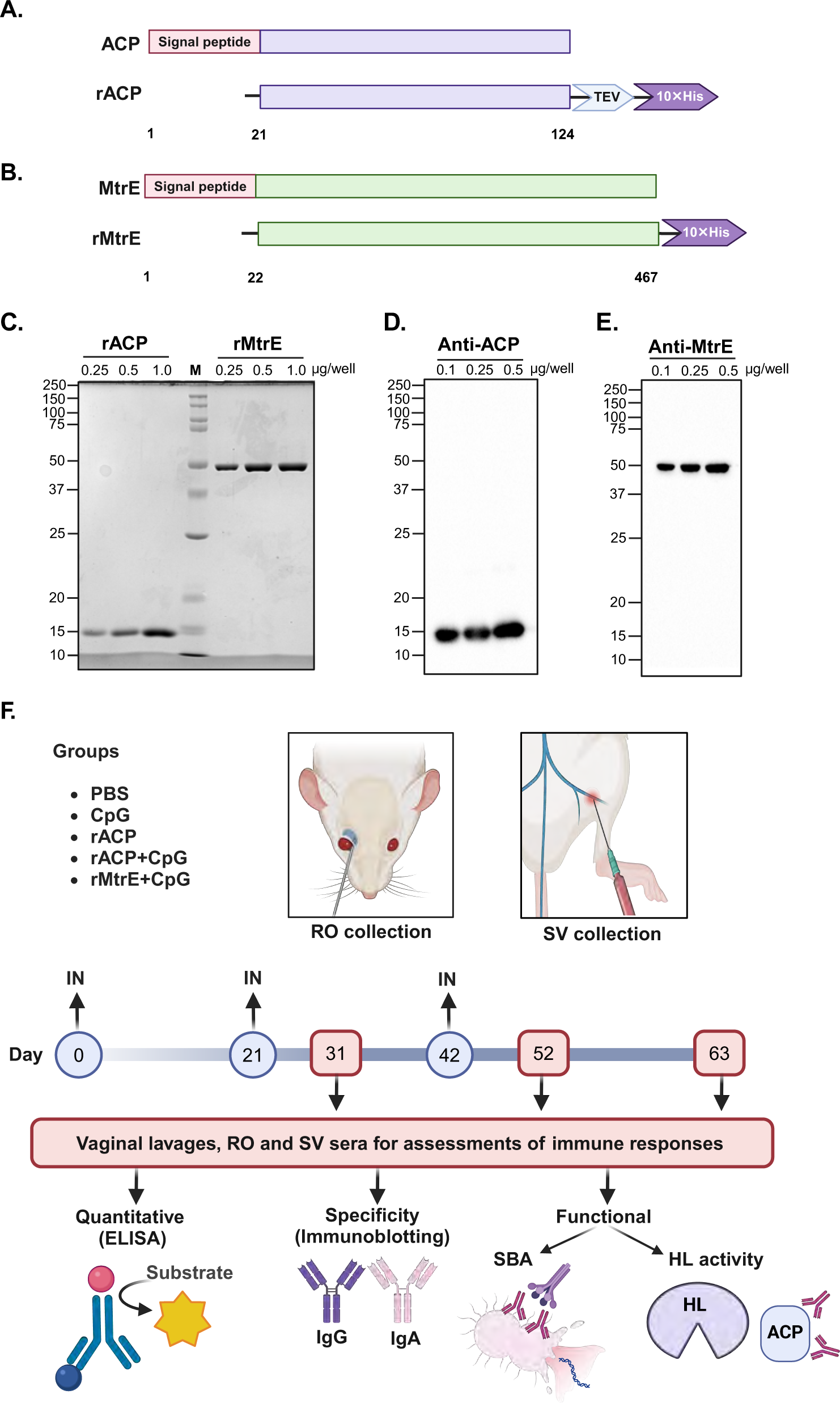
Experimental outline. Panel **A-B**: Schematic of recombinant ACP (rACP) and MtrE (rMtrE) constructs, designed without signal peptides and with C-terminal His-tags for purification (not to scale). **C-E**: rACP (∼11.3 kDa) and rMtrE (∼48.3 kDa) were purified via affinity chromatography and confirmed by SDS-PAGE and Coomassie staining (**C**), and immunoblotting using anti-ACP (**D**) and anti-MtrE (**E**) antisera. Protein amounts are indicated. (**F**) Immunization and sample collection workflow. Female BALB/c mice were randomized into groups (*n*=5–6) and immunized intranasally with PBS, CpG, rACP, rACP+CpG, or rMtrE+CpG (days 0, 21, 42). Serum was collected via retro-orbital (RO) or saphenous vein (SV) bleeds on days 31, 52 and 63 alongside vaginal lavage samples. Immune responses were examined via ELISA, immunoblotting and functional antibody assays including human serum bactericidal assay (SBA) and lysozyme activity (HL).

Recombinant ACP (rACP) was purified as a soluble protein through affinity chromatography, while recombinant MtrE (rMtrE) was extracted from inclusion bodies, affinity purified and refolded to restore its functional conformation. Both antigens were purified to 99% homogeneity and showed migration on SDS-PAGE consistent with their predicted molecular weights of 11.3 kDa for rACP and 48.3 kDa for rMtrE, as confirmed by Coomassie staining (**Fig. 2C**) and immunoblotting with rabbit polyclonal antisera specific to rACP and rMtrE (**Figs. 2D** and **2E**, respectively). Cohorts of mice (*n* = 5–6 per group) were intranasally immunized three times at three-week intervals with one of the following treatments: rACP, rACP+CpG, rMtrE+CpG, CpG alone, or PBS (control), as shown in **Fig. 2F**. The rMtrE antigen was not tested alone due to findings from a recent study [13] identifying CpG1826 as an optimal adjuvant for eliciting a robust Th1-polarized immune response.

To evaluate both mucosal and systemic immune responses, vaginal lavage samples and sera were collected at three time points: days 31, 42, and 63. Blood samples were obtained via retro-orbital (RO) and saphenous vein (SV) routes to assess the specificity of elicited immune responses, antibody levels, and functional antibody activity across different systemic compartments (**Fig. 2F**). This comprehensive experimental design allowed for a robust assessment of the specificity, magnitude, kinetics, and functional quality of immune responses elicited by the immunization regimens. Furthermore, it provided valuable insights into their potential as vaccine candidates while addressing the influence of retro-orbital (RO) and saphenous vein (SV) sampling on specific immunological parameters, such as antibody levels and functional antibody activity.

### Assessment of antigen-specific systemic and mucosal antibody responses elicted by intranasal administration of rACP and rMtrE vaccines

To assess the specificity of immune responses induced by intranasal immunization with mature rACP, rACP+CpG, or MtrE+CpG, terminal pooled sera from RO and SV collections, as well as vaginal lavages, were used in immunoblotting experiments (**Fig. 3**). As expected, no signal was observed in samples containing the isogenic Δ*acp* or Δ*mtrE* mutants or in membranes probed with sera from CpG- or sham-immunized mice. ACP-specific IgG in both RO and SV sera from mice immunized with rACP or rACP+CpG readily cross-reacted with purified rACP and ACP present in the whole-cell lysates of wild-type FA1090 (**Fig. 3A, E**). In contrast, mice administered rACP+CpG but not rACP, generated serum IgA that cross-reacted with rACP, and the signal was consistent across blood collection sites (**Fig. 3B, F**). Notably, while CpG was necessary to induce ACP-specific mucosal IgG (**Fig. 3I**), vaginal IgA was elicited by both ACP-based vaccines (**Fig. 3J**). Further, mice immunized with rMtrE+CpG developed serum and vaginal IgG that specifically recognized both rMtrE and native MtrE in wild-type FA1090 (**Fig. 3C, G** and **K**, respectively), whereas MtrE-specific IgA was detected in vaginal lavages (**Fig. 3L**) but was absent in RO and SV sera collections (**Fig. 3D, H**).

**Figure 3.**
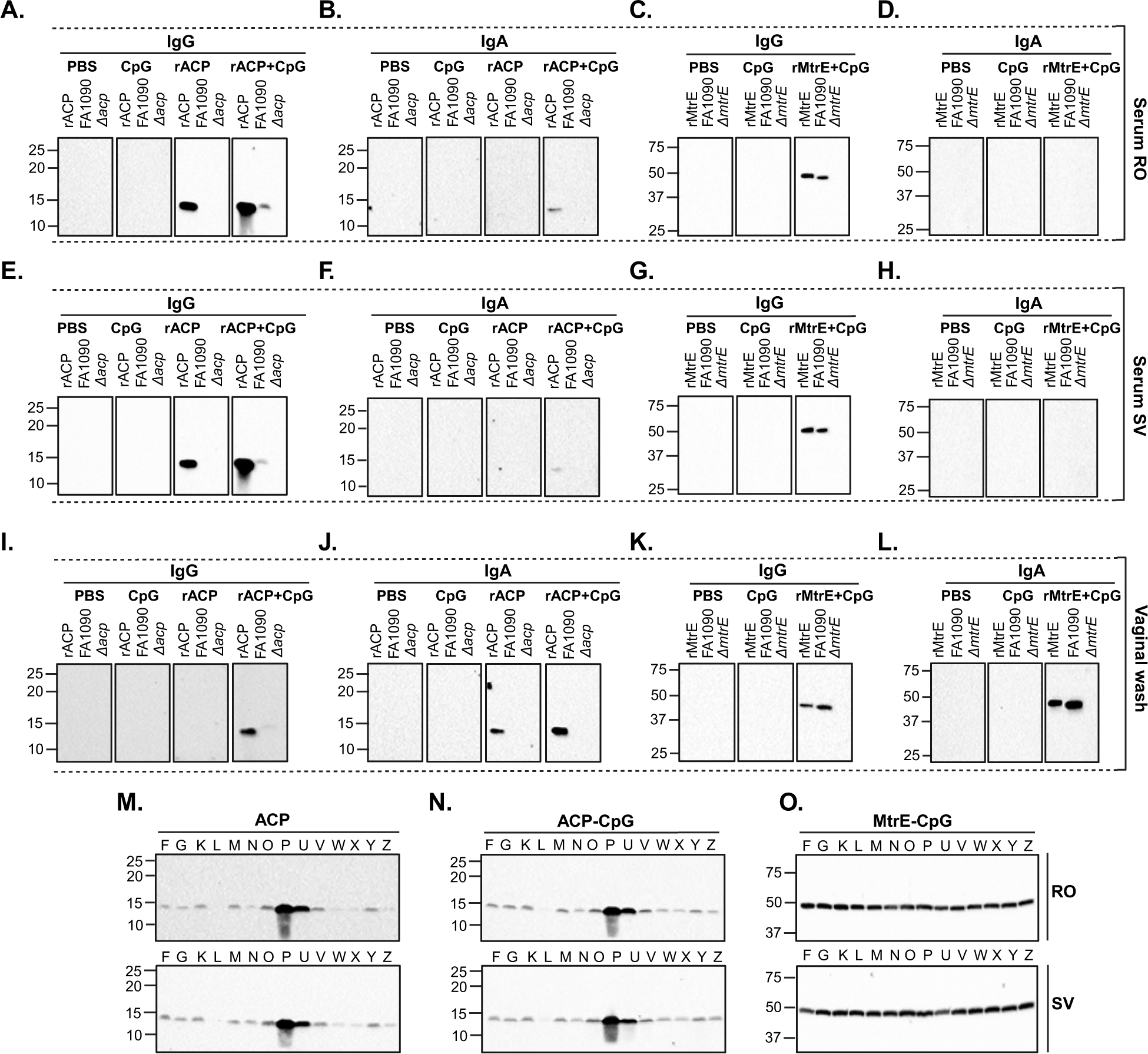
Immunoblotting analysis of ACP- and MtrE-specific IgG and IgA responses. Pooled sera and vaginal washes 63 days post-immunization of female BALB/c mice with recombinant ACP (rACP), rACP combined with CpG (rACP+CpG), recombinant MtrE formulated with CpG (rMtrE+CpG), CpG alone, or PBS were used as primary antibody. Panels **A-H**: IgG (**A**, **C**) and IgA (**B**, **D**) detection in sera collected via retro-orbital (RO; **A-D**) or saphenous vein (SV; **E-H**). Panels **I-L**: IgG (**I**, **K**) and IgA (**J**, **L**) responses in vaginal washes. Panels **M-O**: Comparative immunoblot reactivity of total serum IgG to ACP and MtrE using pooled RO and SV serum samples against whole-cell lysates from 14 genetically and temporally diverse Ng strains in the 2016 WHO Ng reference panel. For all panels, antigen-specific responses are labeled according to experimental conditions. Blots were probed as denoted above, using purified rACP, rMtrE or Ng whole cell lysates.

Together, these results demonstrated that intranasal immunization with rACP or rMtrE-containing vaccines elicited antigen-specific systemic and mucosal IgG responses. Additionally, while the incorporation of CpG with rACP was essential for inducing systemic IgA and vaginal IgG responses, the rACP-based vaccines elicited vaginal IgA independently of CpG. This suggests that rACP alone has intrinsic potential to stimulate localized mucosal immune responses, while CpG is crucial for enhancing systemic and broader mucosal immunity.

### Evaluating the ability of elicited IgG to recognize cognate antigens across diverse

#### N. gonorrhoeae strains

To evaluate the breadth of the antibody responses, we assessed whether serum IgG from mice immunized with rACP, rACP+CpG, or rMtrE+CpG could recognize native ACP or MtrE in whole-cell lysates from 14 genetically and temporally diverse Ng strains included in the 2016 WHO Ng reference panel (**Fig. 3M, N, O**; respectively). Protein bands corresponding to ACP were detected in all tested strains; however, weaker signals were observed in Ng WHO L, N, W, X, and Z, while WHO P and U exhibited the strongest ACP signals across both RO and SV sera collections (**Fig. 3M** and **N**). These differences in ACP detection were not attributable to difference in general protein abundance, as confirmed by Coomassie staining of Ng whole cell lysates (**Fig. S1A**). Additionally, a similar ACP expression pattern was observed when immunoblotting was performed with rabbit anti-ACP serum (**Fig. S1B**), further confirming that the observed differences are not due to protein loading or species-specific variations in antigen recognition. In contrast to the variability seen with ACP, MtrE expression was more uniform across all 14 WHO Ng isolates (**Fig. 3O**). Mice immunized with rMtrE+CpG elicited IgG that consistently recognized MtrE in all tested strains, highlighting its conserved expression.

These results demonstrated that immunization with rACP, rACP+CpG, or rMtrE+CpG vaccines elicited systemic, antigen-specific IgG capable of recognizing ACP or MtrE across genetically diverse Ng strains. No differences in detection levels or cross-reactivity were observed between sera collected via RO and SV routes. While MtrE was uniformly expressed across all isolates, ACP exhibited variability in expression levels, with the highest abundance observed in WHO P and U strain.

#### Quantitative assessments of ACP- or MtrE-reactive antibody types

To quantitatively profile antibody subtypes elicited by immunization with rACP, rACP+CpG, or rMtrE+CpG vaccines, we performed indirect ELISA against purified rACP or rMtrE as coating antigens using RO and SV sera and vaginal lavages collected from individual mice on days 31, 52, and 63 post-initial immunization. The comprehensive analyses included longitudinal kinetic assessments (using AUC log₁₀) of total IgG, IgG1, IgG2a, IgG3, and IgA in RO and SV sera, as well as IgG and IgA in vaginal washes across the three vaccine formulations. Additionally, geometric means of antibody levels between RO and SV samples at the three different time points were determined, along with comparative assessments of serum and vaginal antibody levels elicited by rACP, rACP+CpG, and rMtrE+CpG immunization within each time point (**Figs. 4-7**; **Supplemental Figs. S2-S5**).

**Figure 4.**
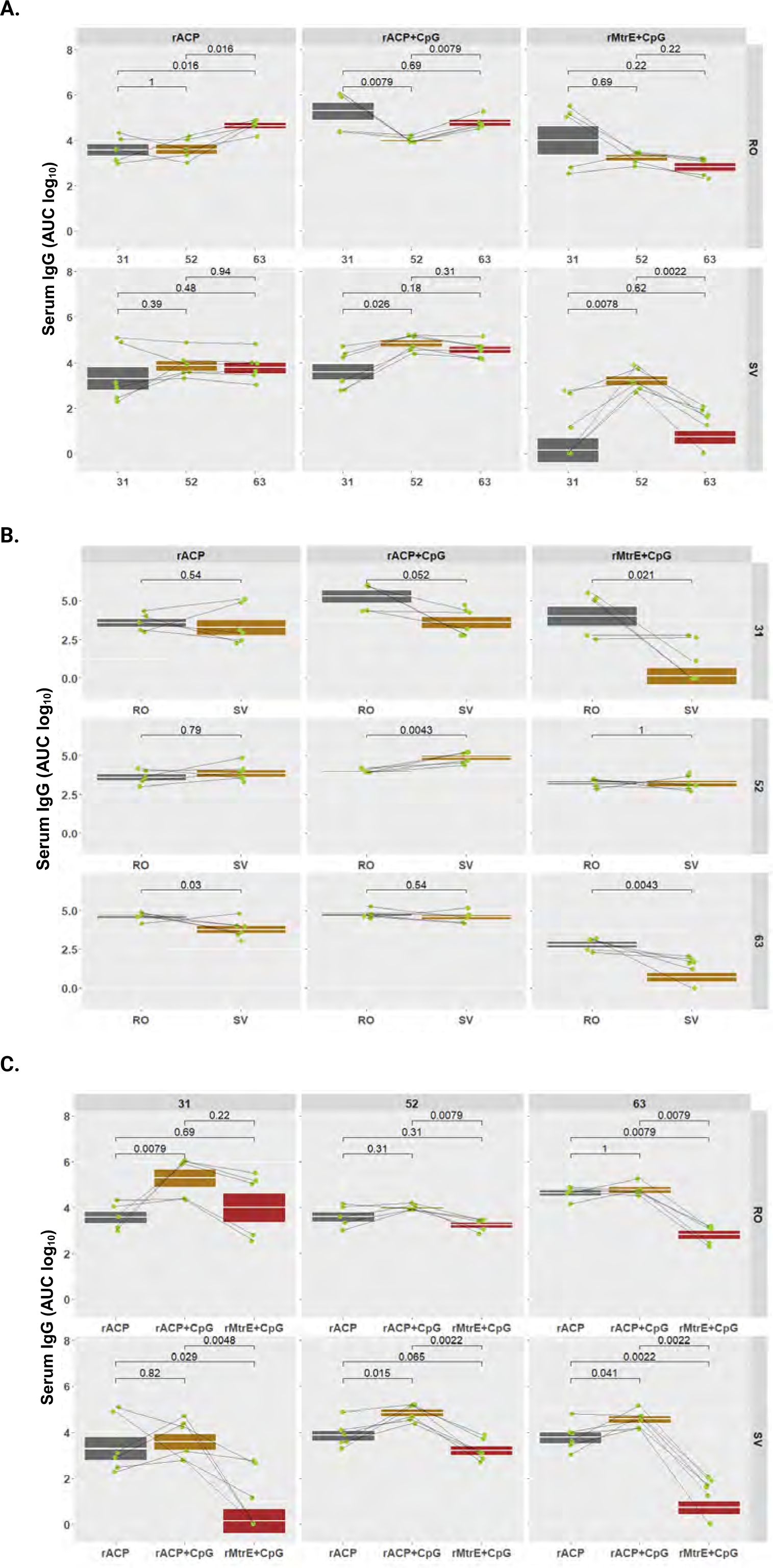
Longitudinal and comparative analysis of serum IgG across antigen formulations and blood collection methods. Serum IgG levels (AUC, log₁₀) were measured in mice immunized intranasally with rACP, rACP+CpG, or rMtrE+CpG and assessed across retro-orbital (RO) and saphenous vein (SV) blood collection sites. Data are presented as geometric means with interquartile ranges, and statistical significance was determined using a two-tailed paired t-test. (**A**) Serum IgG levels over time for RO and SV samples. (**B**) Comparison of serum IgG levels between RO and SV blood collection methods across different vaccine formulations. (**C**) Comparison of IgG responses among different vaccine formulations at different time points within RO and SV groups.

#### Longitudinal and comparative analysis of antibody kinetics across antigen formulations and blood collection methods

Analysis of longitudinal IgG changes revealed that immunization with rACP alone elicited a steady increase in RO samples on day 63 post-immunization *(p*=0.016), whereas the SV group exhibited stable IgG levels with geometric mean AUC log10 of 3.59, 3.86, and 3.77 at days 31, 52 and 63, respectively (**Fig. 4A**, left panel). Formulating rACP with CpG led to early enhancement of IgG followed by a transient suppression at day 52 and an increase at day 63 in RO group, whereas IgG in SV collections increased at day 52 and stabilized in final collections (**Fig. 4A**, middle panel). Mice immunized with rMtrE+CpG demonstrated slightly decreasing IgG levels in RO and a significant peak in SV at day 52 (**Fig. 4A**, right panel). Comparing IgG levels between RO and SV collection sites (**Fig. 4B**), highlighted that RO yielded higher measurements only at day 63 for rACP vaccine (*p*=0.03) and at day 31 and 63 for rMtrE+CpG (*p*=0.021 and *p*=0.0043, respectively). Mice immunized with rACP+CpG vaccine had stable IgG levels and solely on day 52 the SV IgG were higher compared to RO IgG (*p*=0.0043, **Fig. 4B**, middle panel). Comparing IgG levels across antigen formulations at each time point, showed that overall rACP+CpG elicited the strongest response, while IgG induced by rMtrE+CpG was lower compared to ACP-containing vaccines and significantly lower in SV samples (**Fig. 4C**).

IgG1 levels induced by rACP and rACP+CpG vaccines in RO and SV collections were stable over time with a minor decrease for rACP-immunized mice in SV blood on day 52 and a slight but significant incline observed in rACP+CpG RO group consistently across all time points (**Fig. 5A**). In contrast, the rMtrE+CpG-immunized mice showed the most variability in IgG1 levels, particularly within the SV group, with a significant peak on day 52 (*p*=0.0022; **Fig. 5A**, right panel). Comparing the IgG1 levels within each vaccine group across different serum collection methods showed no significant difference except for higher levels for rACP group on day 31 (5.67±0.21 in SV versus 4.38±0.32 in RO, geometric mean±SEM; **Fig. 5B**). Comparing IgG1 levels between rACP-, rACP+CpG-, and rMtrE+CpG-immunized groups at each time point, showed that both rACP and rACP+CpG vaccines yielded similar antibody titers that remained stable over time with a geometric mean from 4.66 to 5.42 (**Fig. 5C**). Mice immunized with rMtrE+CpG had significantly lower IgG1 antibody levels compared to both ACP-containing vaccines on all days consistently in both RO and SV blood collections (**Fig. 5C**).

**Figure 5.**
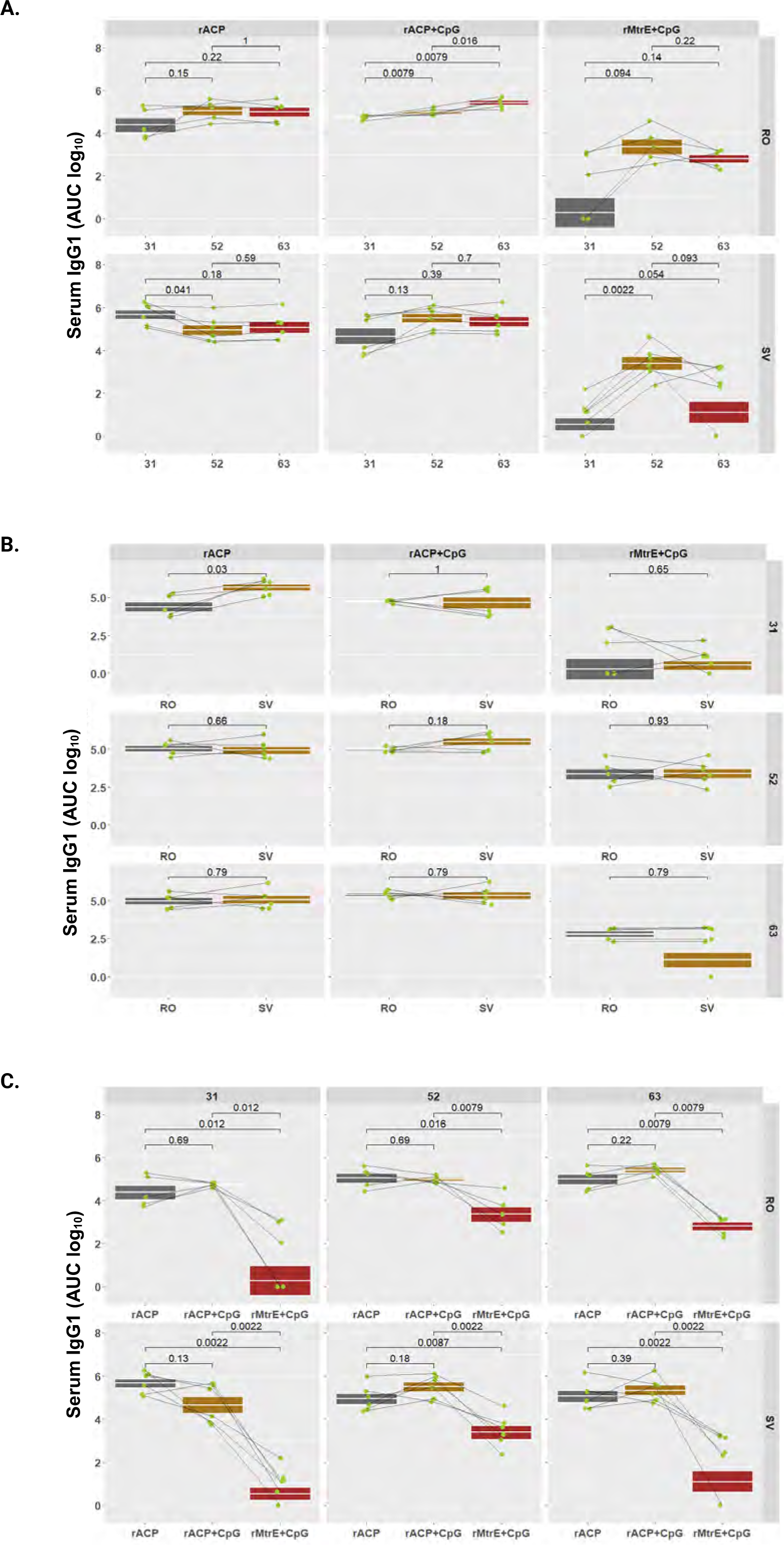
Longitudinal and comparative analysis of serum IgG1. Serum IgG1 levels (AUC, log₁₀) were assessed in mice administered with rACP, rACP+CpG, or rMtrE+CpG and assessed across retro-orbital (RO) and saphenous vein (SV) blood collection sites. Data are presented as geometric means with interquartile ranges, and statistical significance was determined using a two-tailed paired t-test. (**A**) Serum IgG1 levels over time for RO and SV samples. (**B**) Comparison of serum IgG1 levels between RO and SV blood collection methods across different vaccine formulations. (**C**) Comparison of IgG1 responses among different vaccine formulations at different time points within RO and SV groups.

Serum IgG2a levels remained the most stable over time in animals that received the rACP+CpG vaccine, compared to those in the rACP and rMtrE+CpG groups (**Fig. 6A**). In contrast, IgG2a levels in the latter groups showed greater variability, peaking on day 52. However, this increase was significant only for the rMtrE+CpG group in the SV collection compared to days 31 and 63. For all vaccines, both blood collection methods yielded similar IgG2a measurements, except for rMtrE+CpG, which showed significantly higher levels in RO samples compared to SV (*p*=0.046, **Fig. 6B**). As expected, rACP+CpG induced significantly higher IgG2a titers than rACP alone at all time points and across both collection methods (**Fig. 6C**). Similarly, rACP+CpG elicited greater IgG2a levels than rMtrE+CpG at all time points and across all collection sites, except for RO on day 52 (**Fig. 6C**). Further, across all vaccine groups, IgG3 levels remained negligible and comparable to those in PBS- and CpG-immunized animals. However, an exception was observed in rACP+CpG-administered mice within the RO group on day 52, though this response diminished by the terminal collection (**Fig. S2D** and **Fig. S4**). A similar trend was noted for serum IgA, where a significant increase over control groups was detected only in mice immunized with rACP formulated with CpG across all time points with a peak on day 52 post-initial immunization (**Fig. S2E**, **Fig. S5**). This elevation was observed in both RO and SV collections, peaking on day 52 compared to days 31 and 63 (**Fig. S2E**). The increase in serum IgA was significantly greater in the rACP+CpG group compared to the rACP group (*p*=0.011) and the rMtrE+CpG group (*p*=0.097) on day 52 in RO collections. This difference was also observed on days 52 and 63 in SV collections, with statistical significance at *p*=0.0037 for both comparisons on day 52 and *p*=0.0037 and *p*=0.0028 on day 63, respectively (**Fig. S5C**).

**Figure 6.**
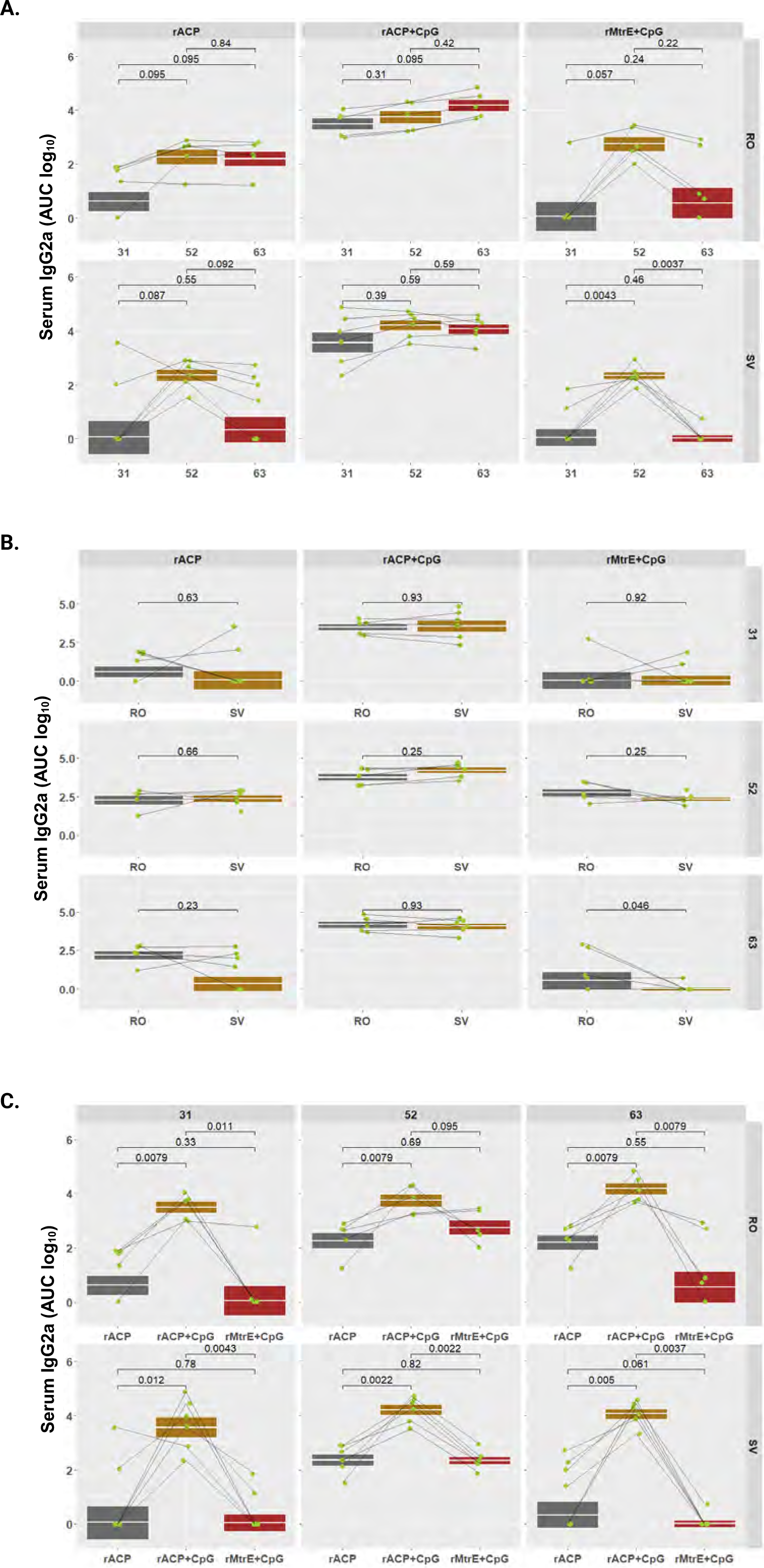
Longitudinal and comparative analysis of serum IgG2a. Serum IgG2a levels (AUC, log₁₀) were assessed in mice administered with rACP, rACP+CpG, or rMtrE+CpG and assessed across retro-orbital (RO) and saphenous vein (SV) blood collection sites. Data are presented as geometric means with interquartile ranges, and statistical significance was determined using a two-tailed paired t-test. (**A**) Serum IgG2a levels over time for RO and SV samples. (**B**) Comparison of serum IgG levels between RO and SV blood collection methods across different vaccine formulations. (**C**) Comparison of IgG2a responses among different vaccine formulations at different time points within RO and SV groups.

**Figure 7.**
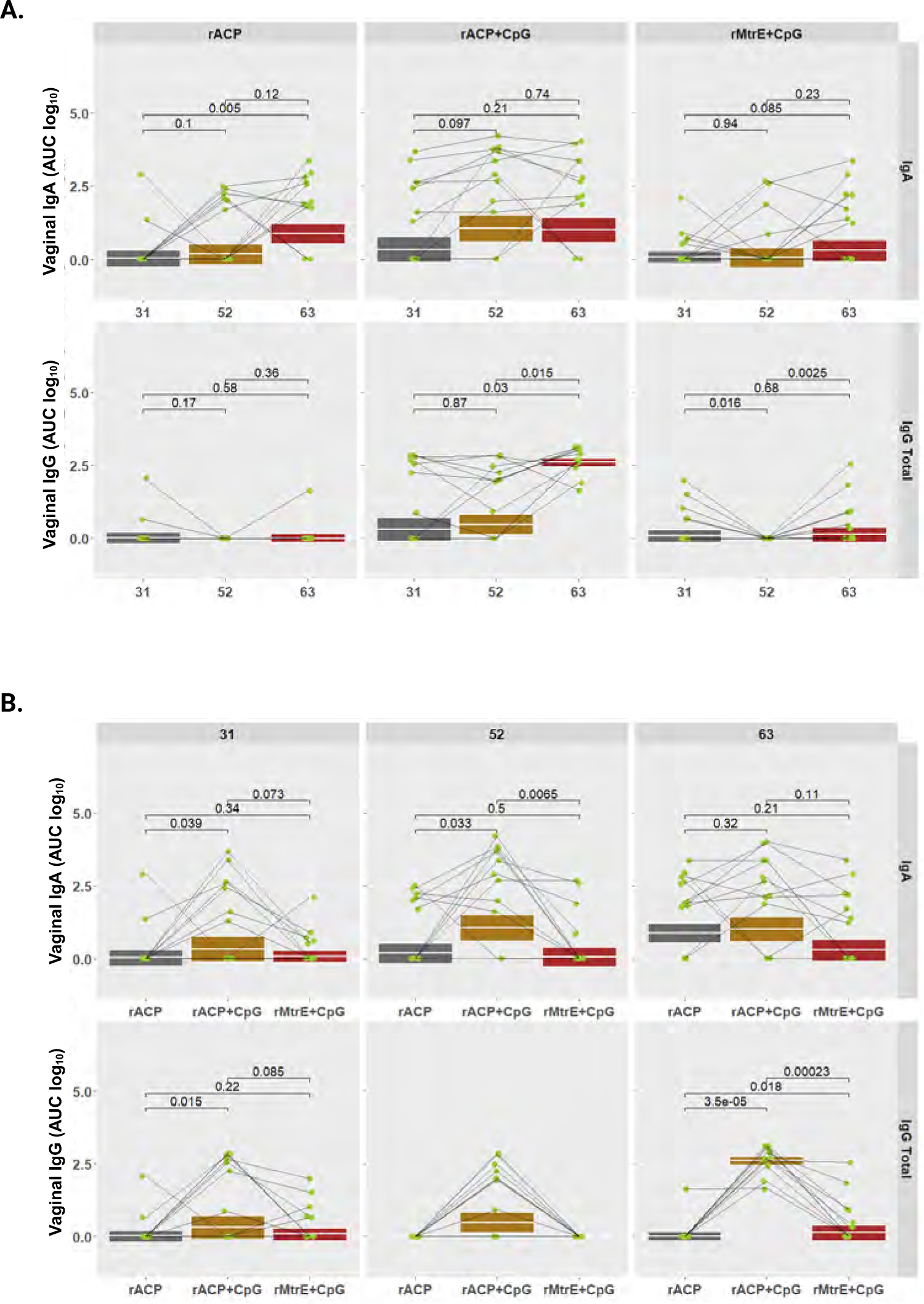
Vaginal mucosal IgA and IgG responses following intranasal immunization with rACP, rACP+CpG, or rMtrE+CpG. (**A**) Kinetics of vaginal IgA (top) and total IgG (bottom) responses (AUC, log₁₀) measured in mice immunized intranasally with rACP, rACP+CpG, or rMtrE+CpG at days 31, 52, and 63 post-immunization. Data are shown for each vaccine group with data from individual mice (green dots) and geometric mean values with interquartile ranges. Statistical comparisons, p-values, between time points within each group are indicated. (**B**) Comparison of vaginal IgA (top) and total IgG (bottom) responses between vaccine formulations (rACP, rACP+CpG, rMtrE+CpG) at each time point (days 31, 52, and 63). Boxplots represent distributions of antibody levels, with significant differences between groups denoted by p-values. Statistical significance was determined using paired t-tests.

Together, these analyses demonstrated that the rACP+CpG formulation elicited the strongest and most stable antibody immune responses, particularly for IgG, IgG2a, and IgA, while rMtrE+CpG induced significantly lower antibody levels across all examined subclasses. Across 45 comparisons of IgG, IgG1, IgG2a, IgG3, and IgA within each vaccine formulation and across different blood collection methods, only eight significant differences were noted: five for IgG (four in RO being higher), one for IgG1 (SV higher), one for IgG2a (SV higher), and one for IgG3 (SV higher). This comparative analysis highlights that RO and SV blood collection methods yield highly comparable antibody kinetic profiles. While SV provided more stable measurements over time, RO detected stronger IgG and IgG2a responses at later time points, particularly for vaccines with weaker immunogenicity, such as rMtrE+CpG.

#### Longitudinal and comparative analysis of vaginal IgG and IgA kinetics across antigen formulations

All three vaccine formulations induced vaginal IgA responses, which increased over time; however, this rise reached statistical significance only in the rACP group on day 63 compared to day 31 (*p*=0.005; **Fig. 7A**). For vaginal IgG, the rACP+CpG vaccine was the most potent, eliciting a steady increase that peaked at day 63, with significantly higher levels compared to days 31 (*p*=0.03) and 52 (*p*=0.015). The geometric mean AUC log10 of vaginal IgG in the rACP group remained stable, showing a slight elevation over control groups at day 31. In contrast, mice immunized with rMtrE+CpG exhibited greater variability in vaginal IgG responses, with a significant decline on day 52 compared to days 31 and 63 (*p*=0.016 and *p*=0.0025, respectively; **Fig. 7B** and **Fig. S 3B**). Compared to rACP alone, rACP+CpG elicited significantly higher IgA levels on days 31 (*p*=0.039) and 52 (*p*=0.033), as well as higher IgG on days 31 (*p*=0.015) and 63 (*p*=0.000035) (**Fig. 7B**). Additionally, rACP+CpG outperformed rMtrE+CpG, inducing significantly higher vaginal IgA on days 31 (*p*=0.073 and 52 (*p*=0.0065), and IgG on day 63 (*p*=0.00023).

These findings demonstrated that rACP alone and rMtrE+CpG primarily induced stronger and more stable vaginal IgA responses compared to IgG over time. Notably, the rACP+CpG vaccine elicited the highest and most durable vaginal IgG and IgA responses, suggesting that this formulation may offer the best long-term antibody protection compared to rACP alone or rMtrE+CpG.

#### Serum bactericidal assay

To assess functional antibody responses, we evaluated pooled terminal antisera from mice immunized with rACP, rACP+CpG, MtrE+CpG, CpG, or PBS using a human serum bactericidal assay (hSBA) against serum-resistant Ng FA1090. All control groups (PBS and CpG alone) exhibited low SBA titers within the 128– 256 range, indicating baseline bactericidal activity. Only CpG-immunized mice showed lower titers in RO samples compared to the SV group and both RO and SV saline controls. Each tested vaccine elicited bactericidal antibodies, achieving ≥50% Ng killing at reciprocal serum dilutions of 2,054—an 8- to 16-fold increase compared to controls (**Table 3**). The hSBA titers for rACP, rACP+CpG, and rMtrE+CpG vaccines were consistently high (2,048) in both retro-orbital (RO) and saphenous vein (SV) samples, suggesting functional antibody responses across antigen-adjuvant formulations. No significant differences in hSBA results were observed between RO and SV blood collection sites in antigen-immunized groups, confirming that hSBA outcomes remain consistent across sampling methods.

**Table 3.**
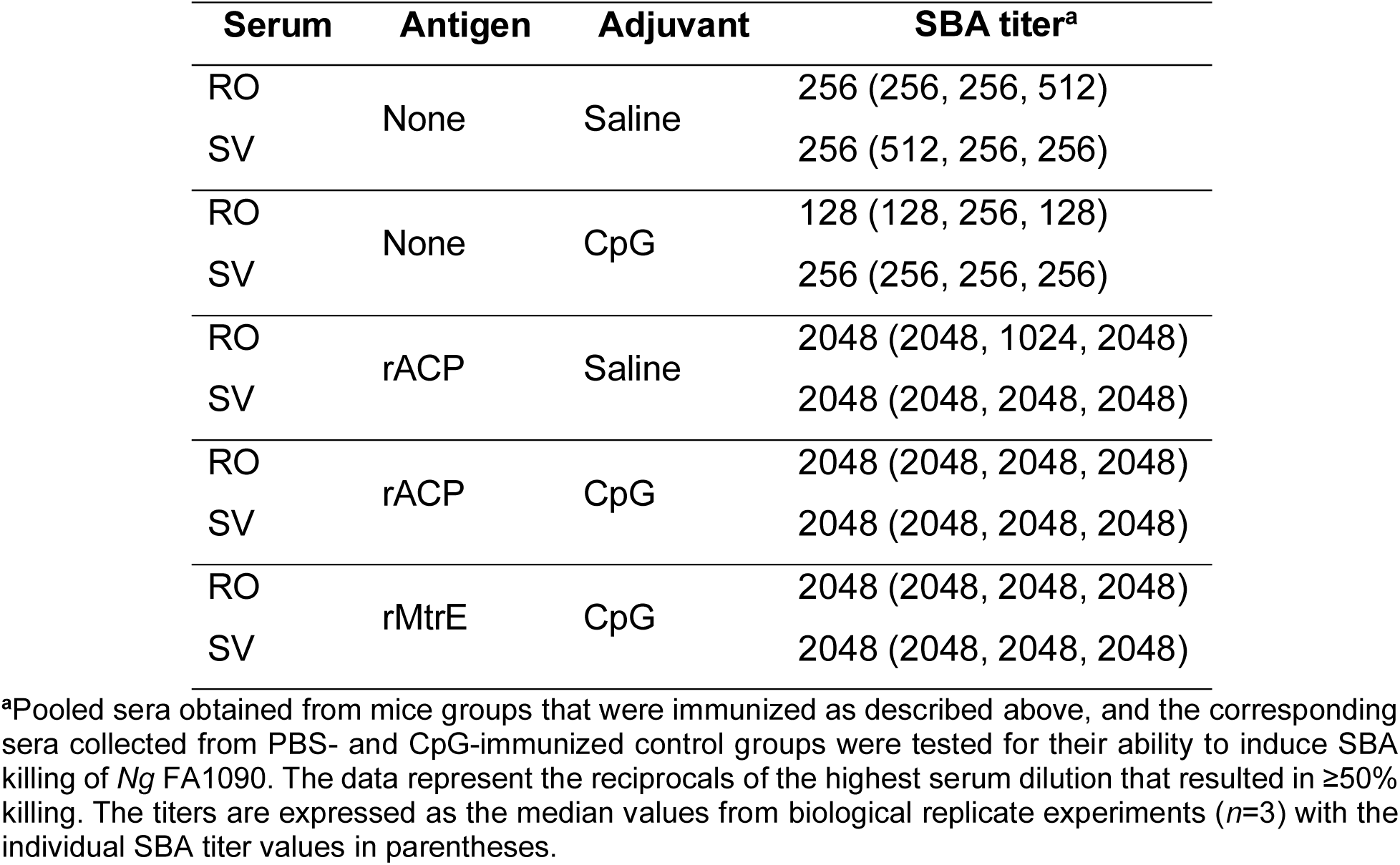
Human complement-dependent serum bactericidal activity of pooled murine RO and SV antisera to rACP, rACP+CpG, or rMtrE+CpG vaccines.

| Serum | Antigen | Adjuvant | SBA titer<sup>a</sup> |
| --- | --- | --- | --- |
| RO | None | Saline | 256 (256, 256, 512) |
| SV |  |  | 256 (512, 256, 256) |
| RO | None | CpG | 128 (128, 256, 128) |
| SV |  |  | 256 (256, 256, 256) |
| RO | rACP | Saline | 2048 (2048, 1024, 2048) |
| SV |  |  | 2048 (2048, 2048, 2048) |
| RO | rACP | CpG | 2048 (2048, 2048, 2048) |
| SV |  |  | 2048 (2048, 2048, 2048) |
| RO | rMtrE | CpG | 2048 (2048, 2048, 2048) |
| SV |  |  | 2048 (2048, 2048, 2048) |

<sup>a</sup>Pooled sera obtained from mice groups that were immunized as described above, and the corresponding sera collected from PBS- and CpG-immunized control groups were tested for their ability to induce SBA killing of *Ng* FA1090. The data represent the reciprocals of the highest serum dilution that resulted in ≥50% killing. The titers are expressed as the median values from biological replicate experiments (*n*=3) with the individual SBA titer values in parentheses.

#### Comparative evaluation of ACP-Induced antibodies in restoring human lysozyme hydrolytic activity across antigen formulations and blood collection methods

Previous studies have shown that Nm and Ng ACP elicit functional blocking antibodies in rabbits, preventing ACP from inhibiting HL-mediated lysis of *Micrococcus lysodeikticus* cell walls *in vitro* [15, 18]. Building on this, we aimed to determine whether mice intranasally immunized with rACP or rACP+CpG produce antibodies capable of inhibiting ACP and whether the choice of blood collection site —(RO) or (SV)—affects antibody levels, activity, and consequently, this interaction. To simulate conditions encountered by Ng in the human host, we used c-type lysozyme purified from human neutrophils and pooled terminal sera from mice immunized with rACP, rACP+CpG, or rMtrE+CpG, as well as control groups. Purified rACP was added, followed by incubation, and the reaction was initiated with fluorescently labeled *M. lysodeikticus* cell walls [19, 29]. As expected, ACP fully inhibited HL hydrolytic activity in the presence of RO and SV sera from mice immunized with PBS, CpG, or rMtrE+CpG (**Fig. 8**). However, antibodies elicited by rACP alone restored 19.3% and 30.05% of HL activity in RO and SV sera, respectively, compared to HL in samples incubated with sera obtained from mice that received PBS. Notably, antisera from rACP+CpG-immunized mice demonstrated a substantial enhancement in HL activity restoration, preventing rACP from inhibiting HL with mean efficacy levels of 59.9% (RO) and 75.2% (SV) compared to mice immunized with rACP alone (*p*<0.001 and *p*<0.0001, respectively).

**Figure 8.**
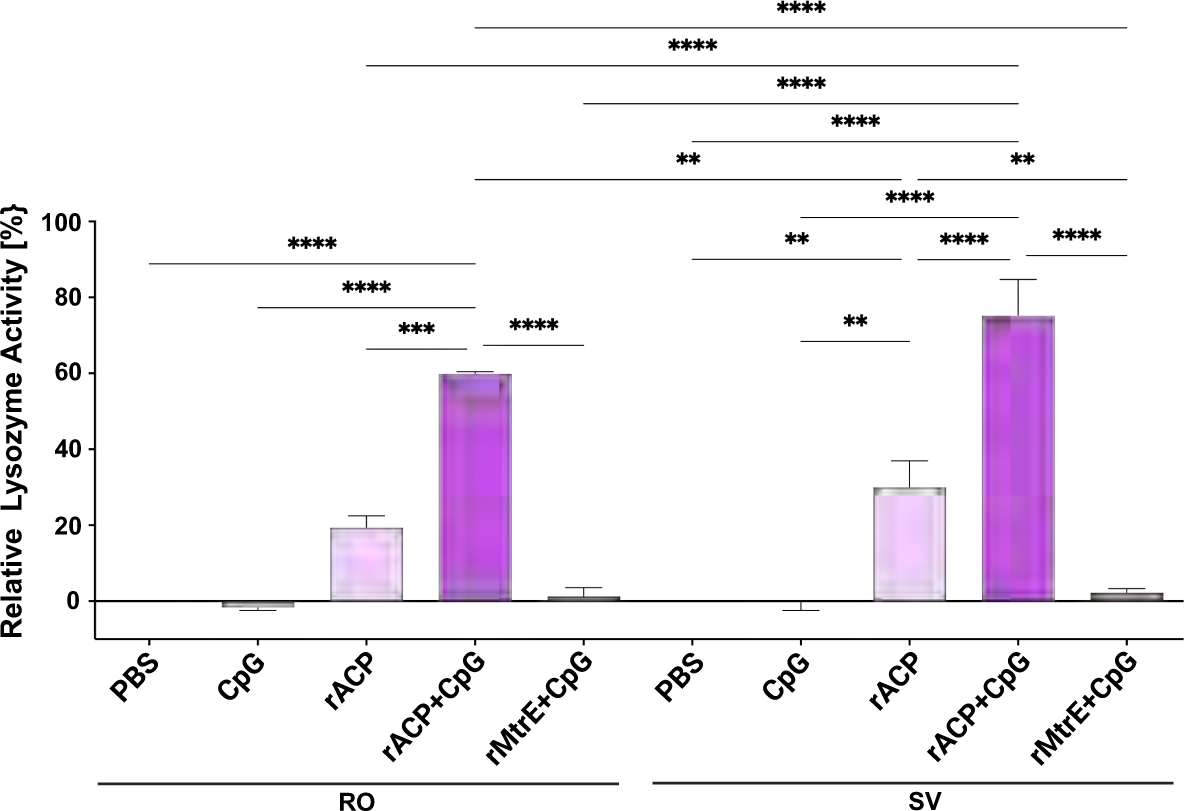
**Mice immunized with rACP-containing vaccines elicit functional blocking antibodies that rescue human c-type lysozyme activity**. Terminal, pooled sera obtained from retroorbital (RO) or saphenous vein (SV) collection sites from mice immunized with PBS, CpG, rACP, rACP+CpG, or rMtrE+CpG were distributed to 96-well black microtiter plates containing human c-type lysozyme (HL). Subsequently rACP was added and after 30 min incubation, the hydrolytic activity of HL was initiated with DQ lysozyme substrate and the fluorescence was measured in a kinetic mode every 2 min for 2 min at 37°C at excitation and emission wavelengths of 485 nm and 530 nm, respectively. The lysozyme activity was calculated according to the standard curve. The percentage of HL activity was computed based on the difference of positive control comprising of HL incubated alone and negative control containing HL and rACP, then normalized to the HL activity in samples incubated with sera obtained from mice that received PBS. Experiments were performed in biological quadruplicates and mean±SEM are presented (\*\**p*<0.01, \*\*\**p*<0.001, \*\*\*\**p*<0.0001).

These findings highlight CpG’s role as a potent adjuvant, amplifying the antibody-mediated restoration of HL activity and countering bacterial immune evasion mechanisms with nearly double the efficacy observed in rACP-alone immunization. Furthermore, the superior functional activity observed in SV blood collections compared to RO collections underscores the impact of blood collection site on immune response assessments.

## DISCUSSION

A critical aspect of this study was evaluating the impact of RO versus SV blood collection on immunological readouts, specifically antibody titers, SBA, and functional blocking antibodies elicited in mice by intranasal delivery of three experimental gonococcal vaccines containing rACP, rACP+CpG, or rMtrE+CpG. To our knowledge, this study represents the first direct comparison of RO and SV blood collection methods in the context of antibody titers and functional antibody responses. Previous studies have examined the effects of different blood sampling techniques on hematological parameters, platelet function, stress markers, and general immune cell profiles [2, 3, 5, 16, 30, 31], but none have specifically assessed their impact on functional antibody levels. Filling this knowledge gap is essential for ensuring methodological consistency in preclinical vaccine studies, as differences in blood collection techniques could influence immunogenicity assessments and immune functional assays.

Our findings demonstrated only eight significant differences across 45 comparisons of total IgG, IgG1, IgG2a, IgG3, and IgA levels across RO and SV blood collection methods. The hSBA titers for rACP, rACP+CpG, and rMtrE+CpG vaccines were consistently high in both RO and SV samples. These results reinforce the robustness of both methods in quantitative antibody assessments and functional hSBA assays. RO blood collection involves disruption of the vascular plexus behind the eye with a capillary tube. The local tissue damage and potential for contamination from glandular tissue that may be disrupted during blood collection could alter analyses performed on collected sera. In addition, RO blood collection occurs under anesthesia which may also impact results. Conversely SV collection may be performed with manual restraint and typically results in minimal tissue trauma. These differences in the blood collection approaches may contribute to some of the minor differences seen in the immunological parameters.

However, antibodies in SV samples exhibited superior restoration of HL hydrolytic activity, suggesting that blood collection methods may differentially impact immunological readouts. The higher HL restoration in SV-derived sera suggests that this method may be more reliable for evaluating functional antibody responses, reinforcing the importance of optimizing blood collection techniques in immunological studies. Furthermore, the use of SV blood collection aligns with the 3Rs, as it is a less invasive method that reduces stress, minimizes risk of ocular trauma, and improves animal welfare, without compromising the integrity of immunological data. Given that many studies in the vaccine field rely on RO blood collection as a standard method, our findings provide compelling evidence for adopting SV blood sampling as a viable and potentially superior alternative, particularly in studies assessing functional immune responses. Standardizing the SV method across vaccine research could enhance data reliability, animal welfare, and procedural refinement, ultimately improving the translational relevance of preclinical vaccine studies. Our *in silico* predictions, coupled with structural mapping of identified B-cell and T-cell epitopes to ACP and MtrE provided novel insights into the antigenic landscape of these vaccine candidates, contributing to the broader efforts in gonococcal vaccine development. In previous studies, both ACP and MtrE were formulated with adjuvants and intraperitoneally inoculated into mice [13, 18]. Human vaccines are commonly administered intramuscularly, subcutaneously, or intranasally [32]. Particularly, for gonococcal vaccine the intranasal administration route may be more attractive considering the mucosal nature of infection [9, 33–35]. In both humans and experimental animals, intranasal vaccine administration effectively induced immune responses in the genital tract and proved more efficient in generating genital antibodies compared to intravaginal inoculation [35–43]. The TLR9 agonist CpG activates human plasmacytoid dendritic cells and B cells, driving a potent Th1-type immune response [44]. Data from several preclinical studies suggest that the Th1-polarized immune response may be important for gonococcal vaccine efficacy. The LOS 2C7 epitope mimic with MPLA or viral replicon particles with PorB enhanced Ng clearance from the murine lower genital tract [45–49]. Intranasally administered experimental vaccines consisting of Ng outer membrane vesicles with microsphere encapsulated interleukin-12 and lipoprotein subunit vaccine rMetQ adjuvanted with cytosine–phosphate–guanine oligodeoxynucleotides (ODN) 1826 class B (CpG B) induced Th1-biased immune responses, mucosal IgG and IgA and resulted in accelerated Ng clearance in mice [33, 34]. Similarly, MtrE and its peptide variants adjuvanted with CpG B, elicited Th1-polarized response, antigen-specific bactericidal antibody responses, and augmented Ng clearance in gonorrhea female mouse model [13]. CpG adjuvant is recognized as a safe and effective intranasal mucosal adjuvant, as no local or systemic inflammation or tissue damage was observed, even at high doses [44, 50]. Our study demonstrated the critical role of CpG as a potent adjuvant in enhancing the immunogenicity and functional efficacy of rACP-based vaccines. CpG significantly amplified antibody-mediated restoration of HL hydrolytic activity, countering bacterial immune evasion mechanisms with efficacy levels nearly double those achieved with rACP alone. It also elicited strong, stable systemic and vaginal mucosal immune responses, reinforcing its potential as an effective mucosal adjuvant [50, 51]. Interestingly, nasal immunization with rACP or rACP+CpG induced higher antibody responses than rMtrE+CpG, suggesting that antigen characteristics influence vaccine performance in the mucosal environment. Despite the advantages of nasal vaccines, challenges remain [44, 52, 53]. One possible explanation for the lower immunogenicity of rMtrE+CpG may be increased mucociliary clearance and/or inefficient uptake of MtrE, which, in its monomeric form, is ∼50 kDa—three times larger than ACP (∼15 kDa) and nearly 10-fold larger in trimeric form. Larger antigens may face barriers in transcellular and paracellular transport, limiting uptake [44, 52]. Future research should focus on optimizing vaccine formulations, including adjuvants that enhance mucosal permeability. Identifying size-dependent parameters for transcellular and paracellular antigen uptake when augmented by permeation enhancers will be critical to improving nasal vaccine efficacy [44, 54]. Such advancements will further refine intranasal gonococcal vaccination strategies, ensuring efficient antigen delivery and robust immune responses.

## MATERIALS AND METHODS

### Epitope mapping

The 3D structures and protein sequences of Ng MtrE (Uniprot ID: Q51006) and ACP (Uniprot ID: A0A379B153) were retrieved from the UniProt Database and visualized using PyMOL v3.1.3. Sequence alignment with homologous proteins was performed using BLAST. B-cell epitopes were predicted with ElliPro (v1.3) [55] identifying epitopes with scores above 0.5, indicative of high surface exposure and immunogenicity. MHC class I epitopes were predicted using NetMHCpan v4.1 [56], targeting 9-14 mer peptides with binding scores above 0.6, suggesting their potential to elicit CD8+ T-cell responses. Similarly, MHC class II epitopes were predicted using NetMHCIIPan v4.1, focusing on 9-14 mer peptides with binding scores above 0.6 for inducing CD4+ T-cell responses. The predicted epitopes were mapped to 3D structures of MtrE and ACP in PyMOL [57].

### Bacterial strains and growth conditions

*Neisseria gonorrheae* (Ng) FA1090 and the 2016 Ng World Health Organization (WHO) reference strains were used in the study [58, 59]. Ng isolates were cultured from frozen stocks onto gonococcal base (GCB, Difco) agar solid medium with Kellogg’s supplements I (1:100) and II (1:1,000) or chocolate agar plates, as indicated, and incubated for 18-20 h at 37°C in a humid 5% CO2 atmosphere [19, 60]. Transparent and non-piliated colonies were sub-cultured onto GCB and used in all experiments. For genetic manipulations in Ng, culture media were supplemented with 40 µg/mL kanamycin and 100 µg/mL streptomycin.

*Escherichia coli* NEB5α was used as the host strain for DNA cloning, whereas *E. coli* BL21(DE3α) and C43(DE3) were used for the overexpression of recombinant ACP (rACP) and recombinant MtrE (rMtrE), respectively. The *E. coli* strains were maintained on Luria-Bertani (LB) agar and broth supplemented with carbenicillin and kanamycin (50 µg/mL).

### DNA manipulations

Primers design and in-silico cloning procedures were performed using genomic DNA sequence of Ng FA1090 and SnapGene (version 2.8, GSL Biotech LLC). Oligonucleotides were synthesized by Integrated DNA Technologies. NEBuilder HiFi DNA Assembly Master Mix, DNA ligase and Q5 High-Fidelity DNA polymerase were procured from New England Biolabs (NEB). Sanger Sequencing was performed at the Center for Quantitative Life Sciences at Oregon State University to verify the final genetic constructs. The *acp* gene was amplified with primers pair ACP Forward (Fwd) 5’GATCCCATGGCCGTCTGAAACACCGGAC3’ and ACP Reverse (Rev) 5’ GATCCTCGAGTGATTAACGTGGGGAACAGTC3’ using FA1090 genomic DNA as a template. The amplified product was digested using *Nco*I/*Xho*I and cloned into similarly digested pET22M [29]. The FA1090 Δ*acp* and the construct for overexpression of rMtrE were described previously [28, 29]. The Δ*mtrE* knockout was generated in the Ng FA1090 according to procedures outlined in prior study using Gibson Assembly [29]. Briefly, to create the DNA block to delete *mtrE*, the upstream and downstream from *mtrE*, and kanamycin resistance cassette were amplified with primer pairs: FwdUpΔ*mtrE* 5’GATCCTTAATTAAGTCTAGAGCATTGGCAAGCGCGCTG3’ and RevUpΔ*mtrE* 5’CGGGACTCTGCCGGCGGTTCTCAGACGG3’, FwdDownΔ*mtrE* 5’GAAGCTAGCTCCGCCCGGGCAAACAAATG3’ and RevDownΔ*mtrE* 5’GCATGCCTGCAGGTTTAAACGATACGGCCTTCCAATACCGAC3’, FwdKan 5’GAACCGCCGGCAGAGTCCCGCTCAGAAG3’ and RevKan5’GCATGCCTGCAGGTTTAAACGATACGGCCTTCCAATACCGAC3’; respectively. Assembled DNA was cloned into pNEB193 and moved into pGCC5 using *Xba*I/*Hinf*I. The pGCC5Δ*mtrE* was linearized and introduced to FA1090 by transformation. The following day, Ng colonies were passaged on selective media with kanamycin and presence of introduced mutation was verified by PCR and immunoblotting with rabbit polyclonal anti-MtrE antisera.

### Antigens expression and purification

Recombinant ACP (rACP) was purified from *E. coli* BL21(DE3) carrying pET22M-rACP grown in 8 L of LB cultures. Bacteria were cultured in an orbital shaker at 37°C, 200 rpm to OD_600_∼0.7. Protein expression was induced by adding 1mM Isopropyl-β-D-1-thiogalactopyranoside (IPTG) for 4 h. The cells were harvested by centrifugation at 4°C, 5,000×g for 15 min. The cell pellets were resuspended in binding buffer (50 mM Na2HPO4, 150 mM NaCl, 10 mM imidazole and 5 % glycerol, pH 7.5) supplemented with 70 µM lysozyme, 0.1µM DNase and protease inhibitor mini tablet (Pierce, 1 tablet/30 mL of buffer). After incubation on ice for 30 min, the cells were lysed using a French Press at 1,500 psi and centrifuged at 12,000×g for 30 min at 4°C to remove the cell debris. The supernatant was filtered using a 0.45 µm nylon membrane. Samples were applied to a HisTrap FF 5 mL Nickel-NTA IMAC column (Cytiva Life Sciences) on an NGC Medium-Pressure Liquid Chromatography system (Bio-Rad) in binding buffer. The bound proteins were washed with a 20% gradient of elution buffer (binding buffer with 250 mM imidazole), collected, pooled and dialyzed against binding buffer using snakeskin tubing (3.5 MWCO) with low stirring overnight at 4°C. The obtained rACP was incubated with Tobacco Etch Virus (TEV) protease supplemented with 1 mM Ethylenediaminetetraacetic acid (EDTA) and 0.5 mM Dithiothreitol (DTT) overnight at 4°C in the ratio of 1:75 to remove the 10His×Tag. Subsequently, the sample was loaded on IMAC to remove TEV and any rACP 10His×Tag. The removal of the 10His×Tag was confirmed by immunoblotting as described below. The rACP was concentrated using Vivaspin 20 centrifugal concentrator and the protein concentrations was quantified by Qubit Protein Assay using Qubit 4 Fluorometer (Invitrogen). The purity of the protein was determined by SDS-PAGE and Western blotting.

To purify rMtrE, *E. coli* C43(DE3) carrying pT7-7K MtrE was grown as described [28]. Bacteria were harvested by centrifugation as outlined above for rACP, cell pellet was resuspended in lysis buffer containing 50 mM NaH2PO4, 300 mM NaCl, 10 mM imidazole, pH 8.0 supplemented with 70 µM lysozyme, 0.1 µM DNase and protease inhibitor mini tablet (Pierce) per 30 mL buffer. Cells were disrupted by sonication 12×10 sec pulse with 30 sec pause. The cell lysate was centrifuged at 10,000×g for 30 min at 4°C and the inclusion bodies were washed twice and resuspended in 20 mM Tris-HCl, pH 8.0. After a brief sonication, the inclusion bodies were pelleted by centrifugation at 12,000×g for 20 min at 4°C, suspended in freshly made denaturation buffer containing 8 M Urea, 100 mM NaH2PO4, 10 mM Tris, pH 8.0 and incubated with shaking for 30 min. The solubilized fraction was subjected to centrifugation at 10,000×g for 10 min at RT. The supernatant was mixed with equilibrated Ni-NTA agarose (Qiagen) for 30 min at room temperature. Mixture was loaded into a BioRad column, and the settled resin was washed with five bed volumes of 6 M Urea, 100 mM NaH2PO4, 10 mM Tris, pH 8.0. The denatured rMtrE was refolded on the resin by washing with five bed volumes of decreasing urea gradient. The refolded rMtrE was eluted with lysis buffer containing 250 mM imidazole, then dialyzed in PBS overnight. The protein concentration was quantified by the DC Protein Assay (BioRad) and the purity of the samples was analyzed by SDS-PAGE and immunoblotting.

### Immunization studies

Immunization studies were conducted using female BALB/c mice (4-6 weeks old, Charles River Laboratories, NCI BALB/c strain). Mice (*n*=11/group) were immunized three times in three-week intervals intranasally (IN; d0, d21, d41) with rACP (2.6 µg/µL), rACP adjuvanted with CpG (ODN2395; Invivogen), or rMtrE (3.3 µg/µL) adjuvanted with CpG, whereas the control groups received PBS or CpG (at 10 µg/dose; total volume 10 µL; 5 µL per nostril). Retro-orbital and saphenous blood collections, and vaginal washes were performed ten days post booster doses (d31 and d52) and the final collections were performed on d63. Each immunization group was split into two mice cohorts: RO bleeding (*n*=5) and SV sampling (*n*=6). RO blood collections and vaginal washes were performed on anesthetized mice (2.0% isoflurane, 1L/m O_2_). Saphenous blood was collected from restrained mice with the hindlimb extended. Briefly, a microcapillary tube was introduced into the medial canthus at a 45° caudomedial angle to collect blood from retroorbital sinus. For SV blood collection, the medial saphenous was visualized and punctured with a 25-gauge needle. A small amount of ophthalmic ointment was applied to smooth the fur, preventing blood absorption by the hair and allowing the drop to pool for collection in a capillary tube. Blood and vaginal washes were centrifuged at 3,000×g for 10 min at 4°C and serum and vaginal lavage supernatants were stored at -80°C until further use.

### Ethics Statement

Animal experiments were performed according to the guidelines of the Association for the Assessment and Accreditation of Laboratory Animal Care under a protocol #IACUC2021-0244, which were approved by the Oregon State University’s Institutional Animal Care and Use Committee. The animal facilities meet the housing service and surgical standards set forth in the “Guide for the Care and Use of Laboratory Animals” NIH Publication No. 85-23. Animals are maintained under the supervision of a full-time veterinarian. For all experiments, mice were humanely euthanized upon reaching the study endpoint using a compressed CO_2_ gas cylinder, as per the institution guidelines, which follow those established by the 2020 American Veterinary Medical Association Panel on Euthanasia (https://www.avma.org/resources/pet-owners/petcare/euthanasia).

### SDS-PAGE and Immunoblotting

The Ng whole cell lysates were prepared by suspending in Laemmli Sodium dodecyl sulfate (SDS) buffer and standardized by OD_600_; recombinant protein samples were loaded based on protein concentration. The samples were resolved on 4%–15% Bio-Rad Criterion TGX (Bio-Rad) and visualized by Colloidal Coomassie staining or transferred onto 0.2 µM nitrocellulose membranes (Trans-Blot Turbo, Bio-Rad) using a Trans-Blot Turbo transfer system (Bio-Rad). The membranes were blocked using 5% non-fat milk in PBST for 1 h at room temperature and probed overnight with pooled sera (1:5,000) or vaginal lavages (1:50) from the terminal collection (D63). After washing with PBST, the immunoblots were probed with HRP conjugated goat anti-mouse IgG or IgA at 1:10,000 dilution (Southern Biotech) and developed using ClarityTM ECL substrate (BioRad). The stained gels and developed immunoblots were visualized in ChemiDocTM MP imaging system (BioRad).

### Enzyme-Linked Immunosorbent Assays

For Enzyme-Linked Immunosorbent Assays (ELISA) U-shaped high-binding 96-well plates were coated with purified rACP, or rMtrE, (100 ng/well) diluted in coating buffer (50 mM Na2CO3-NaHCO3, pH 9.6) overnight at 4°C. The plates were washed with PBS supplemented with 0.05% Tween-20 (PBST) using BioTek 405 LS Microplate washer (Agilent Technologies). Subsequently, Block Ace (Bio-Rad) dissolved in PBST was added for an hour at 37°C. Serial dilutions of sera (starting dilution of 1:729) or vaginal lavages (starting dilution of 1:27) were made on Deepwell sample plates and transferred to corresponding coated plates using Viaflo 96 channel pipette (Integra Biosciences) and incubated for an hour. Following three washes with PBST, the plates were incubated with HRP conjugated goat anti-mouse antisera (dilution 1:10,000) specific for total IgG, IgG1, IgG2a, IgG3, and IgA (Southern Biotech). After another wash, the reactions were developed using 1-Step UltraTM 3,3’,5,5’-tetramethylbenzidine (TMB) substrate (ThermoScientific). The absorbance values for all dilutions were corrected by subtracting the average blank absorbance. These corrected values were used to calculate the area under the curve (AUC), with the baseline set as the average blank absorbance. The areas of the positive peaks were logarithmically transformed and plotted using the ggplot2 package in R.

### Serum Bactericidal Assays

The human complement dependent serum bactericidal assay (SBA) was performed according to method previously described [19]. Briefly, heat inactivated murine sera from immunized and control groups were serially two-fold diluted in Hanks’ Balanced Salt Solution (HBSS) starting from 1:64 to 1:2048. Non-piliated Ng FA1090 grown on chocolate agar plates was harvested and resuspended in HBSS to an OD600 of 0.1 (∼1×10^8^ CFU/mL). Aliquots of 1×10^3^ bacteria in 40 µL of HBSS were added to wells containing diluted test sera. After 15 min, 10 µL of either IgG/IgM-depleted normal human serum (NHS) or heat-inactivated NHS (HI-NHS) was added at 10% (v/v) complement source and incubated for an hour. Finally, 5 µL of aliquots and their corresponding 10-fold dilutions were spot inoculated onto chocolate agar plates and allowed to grow overnight for CFU calculations. Ng with NHS, Ng with HI-NHS, Ng with test sera and HI-NHS, and Ng alone served as controls. The average percent killing was enumerated by comparing the number of CFUs recovered from Ng incubated with test sera and NHS to the number of CFUs recovered from Ng incubated with experimental sera and HI-NHS in three independent studies. The SBA titers values were determined as the those that achieved 50% Ng killing.

### Assessment of lysozyme activity

To determine whether immunization with rACP and rACP-CpG generates functional antibodies that can inhibit rACP’s activity as a lysozyme inhibitor, we used the EnzChek Lysozyme Assay Kit (ThermoFisher) with the following modifications. A lysozyme standard curve of 0, 100, 200, 300, and 400 U/mL lysozyme was included in each assay. To mimic the physiological conditions in the human host, a c-type lysozyme purified from human neutrophils (HL, Sigma) was chosen for measuring rACP-dependent lysozyme inhibition. HL and terminal, pooled sera from immunized or control groups were mixed in a reaction buffer containing 0.1 M sodium phosphate pH 7.5, 0.1 M sodium chloride and 2 mM sodium azide. In a black 96-well plate (Costar), 8 µL serum sample was mixed with 40.5 µL of master mix containing 38.5 µL of reaction buffer and 2 µL HL (10 µM) in each well. Subsequently, 1.5 µL rACP (20 µM) was added to each well and mixed carefully. After 30 min incubation at 37°C, the reaction was initiated by addition of 50 µL DQ lysozyme substrate (50 µg/mL) per well and the fluorescence was immediately measured in a kinetic mode every two min for 20 min at 37°C using a BioTek Synergy HT Microplate Reader at excitation and emission wavelengths of 485 nm and 530 nm, respectively. The lysozyme activity was calculated according to the standard curve. The percentage of HL activity was computed based on the difference of positive control comprising of HL incubated alone (0.4 µM) and negative control containing HL and rACP (0.4 + 0.6 µM; respectively), then normalized to the HL activity in samples incubated with sera obtained from mice that received PBS. Experiments were performed in biological quadruplicates and mean±SEM are presented.

### Statistical analyses

For the ELISA analysis, the geometric mean and standard error of the mean (SEM) were calculated for the AUC across five immunoglobulin classes or subclasses of antibodies in serum (total IgG, IgG1, IgG2a, IgG3, and IgA) and two immunoglobulin classes in vaginal lavage (total IgG and IgA). Pairwise comparisons were conducted using t-tests for different antibody types, serum collection methods, vaccine groups, and submental blood collections. All plots were generated using the ggplot2 package in R. In experiments assessing lysozyme activity, mean and SEM from biological quadruplicate experiments were calculated, and statistical difference were compared using one-way ANOVA.

### Visualization and illustrations

Graph for lysozyme assay was generated using GraphPadPrism10 and illustrations were created using BioRender.com to visually represent the study findings.

## SUPPLEMENTARY INFORMATION

Supplementary materials are included as PDF file.

## AUTHOR CONTRIBUTIONS

**AES** contributed to conceptualization, methodology, data curation, writing—original draft preparation, writing—review & editing, supervision, project administration, and funding acquisition. **AC** contributed to methodology, validation, formal analysis, investigation, data curation, writing—original draft preparation, writing—review & editing, and visualization. **YS** contributed to methodology, validation, formal analysis, investigation, and writing methods section and review of final manuscript. **JN** contributed to methodology, validation, formal analysis, investigation, writing—review & editing, and visualization. **CL** contributed to methodology, validation, formal analysis, investigation, writing—review & editing, and visualization. **ACh** contributed to methodology, validation, formal analysis, investigation, and review. **JS** contributed to methodology and writing—review & editing. All authors read and approved the final manuscript.

## DATA AVAILABILITY STATEMENT

The raw data supporting the conclusions of this article will be made available by the authors on request.

