## Supplemental Figures for "Bridging Gaps in Antibody Responses and Animal Welfare: Assessing Blood Collection Methods and Vaginal Immunity in Mice Immunized with Intranasal Gonococcal Vaccines"

### Supplementary Figures

**Supplementary Figure S1. Expression of ACP across *Neisseria gonorrhoeae* reference strains. (A**) SDS-PAGE analysis of total protein lysates from *Neisseria gonorrhoeae* FA1090, isogenic *Δacp* mutant, and the 2016 WHO Ng reference strains (F, G, K, L, M, N, O, P, U, V, W, X, Y, Z). Purified recombinant ACP (rACP) was included as a control. (**B**) Western blot analysis with polyclonal rabbit serum against rACP detecting ACP expression in the same Ng whole cell lysates as in panel (A). rACP was used as a positive control. The *Δacp* mutant serves as a negative control, confirming specificity. Molecular weight markers (kDa) are indicated on the left.

**Supplementary Figure S2. Serum antibody responses following intranasal immunization with rACP, rACP+CpG, or rMtrE+CpG.** Serum antibody levels (AUC, $\log_{10}$) were measured in mice immunized intranasally with CpG, PBS, rACP, rACP+CpG, or rMtrE+CpG and assessed across retro-orbital (RO) and saphenous vein (SV) blood collection sites at days 31, 52, and 63 post-initial immunization. Data are presented as geometric means with interquartile ranges. Data are presented as log-transformed antibody titers, with individual data points from each mouse (green dots) and boxplots representing geometric mean values with interquartile ranges. (**A**) Total serum IgG titers, (**B**) Serum IgG1 titers, (**C**) Serum IgG2a titers, (**D**) Serum IgG3 titers, and (**E**) Serum IgA titers.

**Supplementary Figure S3. Vaignal antibody responses following intranasal immunization with rACP, rACP+CpG, or rMtrE+CpG.** Vaginal antibody levels (AUC, $\log_{10}$) were measured in mice immunized intranasally with CpG, PBS, rACP, rACP+CpG, or rMtrE+CpG and assessed at days 31, 52, and 63 post-initial immunization. Data are presented as geometric means with interquartile ranges. Data are presented as log-transformed antibody titers, with individual data points from each mouse (green dots) and boxplots representing geometric mean values with interquartile ranges. (**A**) Total vaginal IgG titers, (**B**) Vaginal IgA titers.

**Supplementary Figure S4. Longitudinal and comparative analysis of serum IgG3 across antigen formulations and blood collection methods.** Serum IgG3 levels (AUC, $\log_{10}$) were measured in mice immunized intranasally with rACP, rACP+CpG, or rMtrE+CpG and assessed across retro-orbital (RO) and saphenous vein (SV) blood collection sites. Data are presented as geometric means with interquartile ranges, and statistical significance was determined using a two-tailed paired t-test. (A) Serum IgG3 levels over time for RO and SV samples. (B) Comparison of serum IgG3 levels between RO and SV blood collection methods across different vaccine formulations. (C) Comparison of IgG3 responses among different vaccine formulations at different time points within RO and SV groups.

**Supplementary Figure S5. Longitudinal and comparative analysis of serum IgA.** Serum IgA levels (AUC, $\log_{10}$) were assessed in mice administered with rACP, rACP+CpG, or rMtrE+CpG and assessed across retro-orbital (RO) and saphenous vein

(SV) blood collection sites. Data are presented as geometric means with interquartile ranges, and statistical significance was determined using a two-tailed paired t-test. (**A**) Serum IgA levels over time for RO and SV samples. (**B**) Comparison of serum IgA levels between RO and SV blood collection methods across different vaccine formulations. (**C**) Comparison of IgA responses among different vaccine formulations at different time points within RO and SV groups.

### Supplementary Fig. 1

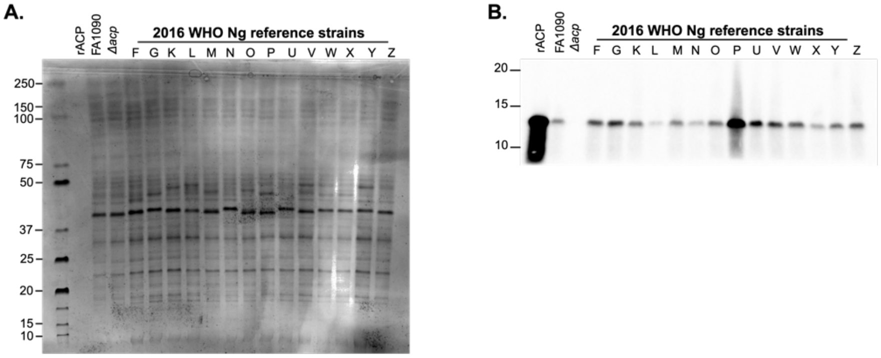

### Supplementary Fig. S2

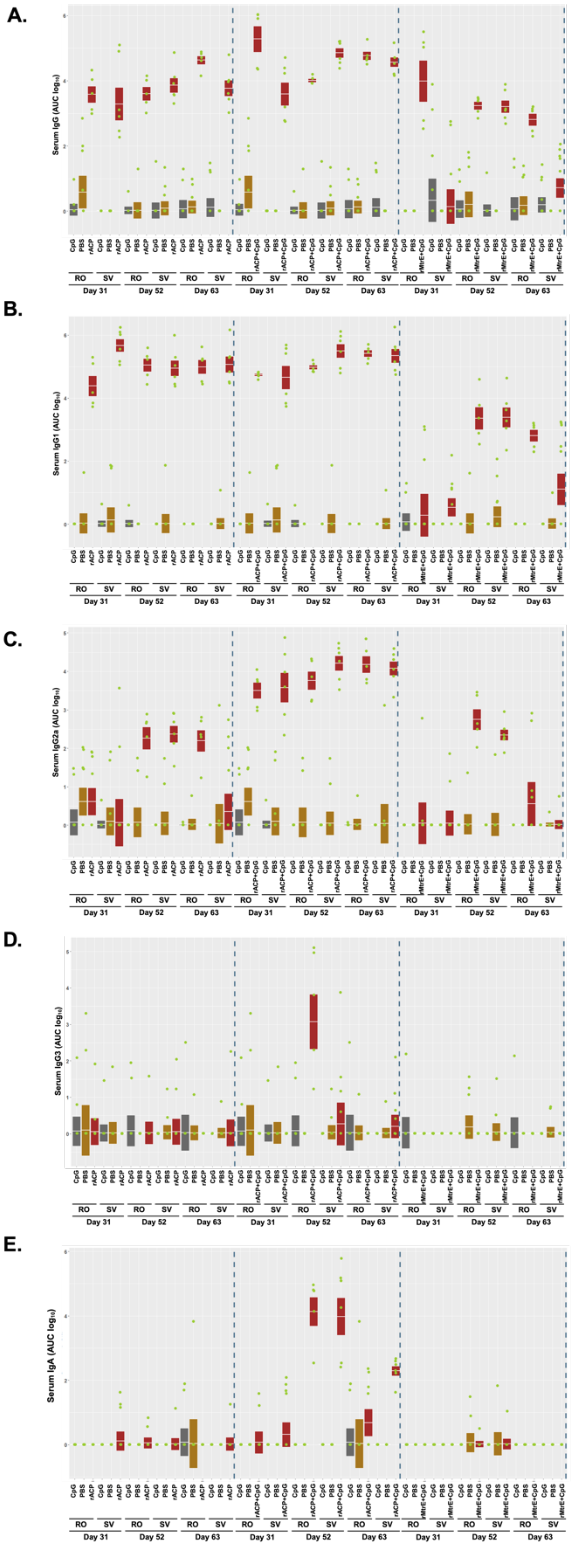

**Supplementary Fig. S3**

**A.**

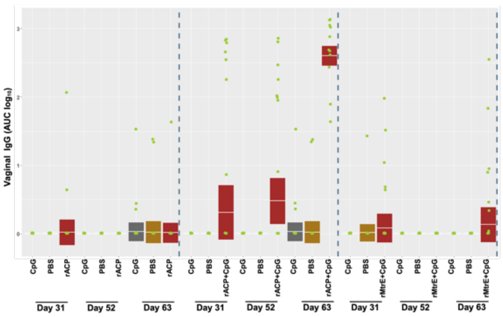

**B.**

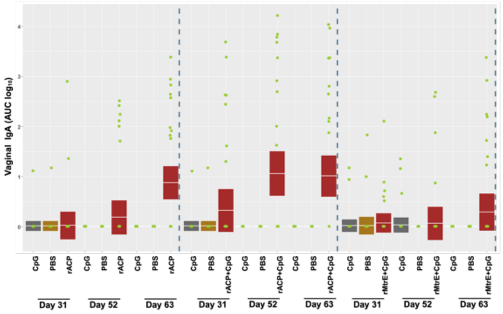

**Supplementary Fig. S4**

**A.**

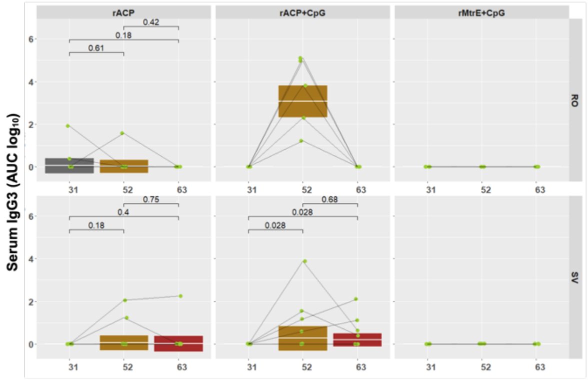


**B.**

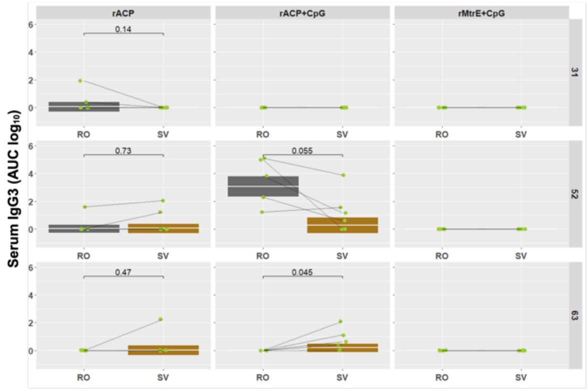


**C.**

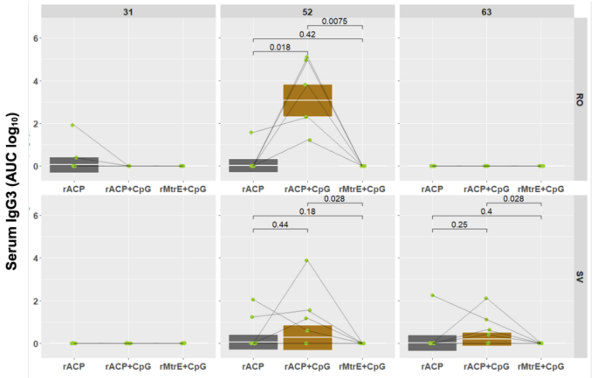

**Supplementary Fig. S5**

**A.**

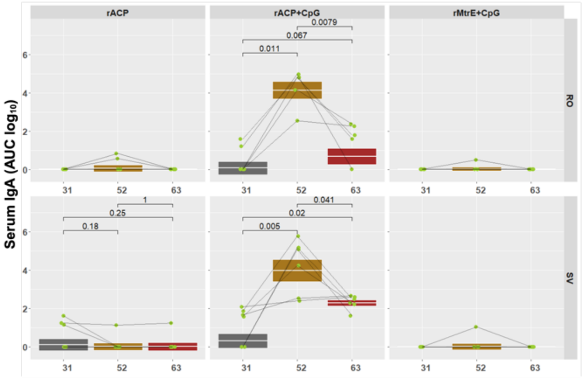


**B.**

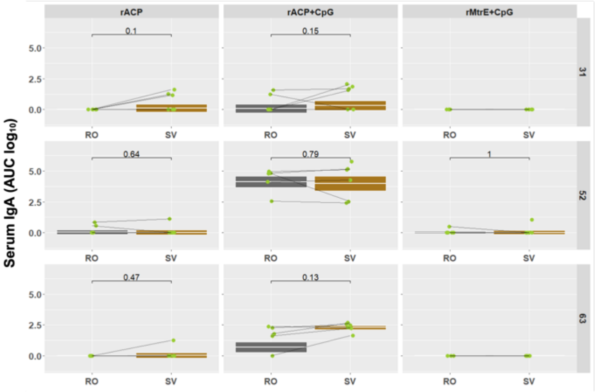


**C.**

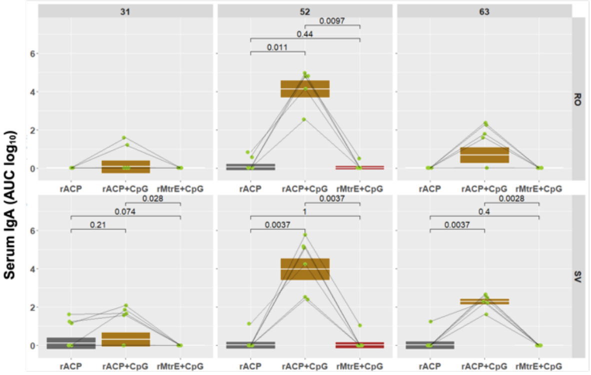
